# The SAGE genetic toolkit enables highly efficient, iterative site-specific genome engineering in bacteria

**DOI:** 10.1101/2020.06.28.176339

**Authors:** Joshua R. Elmore, Gara N. Dexter, Ryan Francis, Lauren Riley, Jay Huenemann, Henri Baldino, Adam M. Guss, Robert Egbert

**Author notes:** Author Contributions (CRediT taxonomy) Conceptualization – JRE, AMG, RE. Methodology – JRE, GND, LR, AMG. Validation – RF, HB. Investigation – JRE, GND, RF, JH, HB. Writing – original draft preparation – JRE. Writing – review and editing – JRE, AMG, RE. Project Administration – JRE, AMG, RE. Funding Acquisition – JRE, AMG, RE.

## Abstract

Sustainable enhancements to crop productivity and increased resilience to adverse conditions are critical for modern agriculture, and application of plant growth promoting rhizobacteria (PGPR) is a promising method to achieve these goals. However, many desirable PGPR traits are highly regulated in their native microbe, limited to certain plant rhizospheres, or insufficiently active for agricultural purposes. Synthetic biology can address these limitations, but its application is limited by availability of appropriate tools for sophisticated, high-throughput genome engineering that function in environments where selection for DNA maintenance is impractical. Here we present an orthogonal, <u>S</u>erine-integrase <u>A</u>ssisted <u>G</u>enome <u>E</u>ngineering (SAGE) system, which enables iterative, site-specific integration of up to 10 different DNA constructs at efficiency on par or better than replicating plasmids. SAGE does not require use of replicating plasmids to deliver recombination machinery, and employs a secondary serine-integrase to excise and recycle selection markers. Furthermore, unlike the widely utilized pBBR1 origin, DNA transformed using SAGE is stable without selection. We highlight SAGE’s utility by constructing a 287-member constitutive promoter library with a ∼40,000-fold dynamic range in *P. fluorescens* SBW25. We show that SAGE functions robustly in diverse α- and γ-proteobacteria, thus providing evidence that it will be broadly useful for engineering industrial or environmental bacteria.

## Introduction

Sustainable enhancements to crop productivity and increased resilience to adverse conditions, such as the burgeoning stresses on ecosystems from climate change, are critical for both food security and biofuel production^3^. The application of plant growth promoting rhizobacteria (PGPR) in agriculture has the promise to both enhance resistance to adverse conditions (*e.g.* drought^4^, high salinity^5^, phytopathogen infection^6^, pest infestation^7^), and reduce reliance on agrochemicals^8^ - thus ameliorating the negative environmental impact of current agriculture methods^1,2^. However, many desirable PGPR functions (*e.g.* nitrogen fixation, pathogen resistance, phosphate solubilization) are complexly regulated in their native microbe^9,10^, found only in microbes with narrow plant host-range, or insufficiently active for agricultural purposes. One solution to is combine synthetic biology with environmental microbiology to enhance desirable traits in PGPR, to transfer novel PGP functionalities between bacteria that thrive in distinct ecosystems, and to enable PGPR association with new host plants.

However, the application of synthetic biology with PGPR is limited by the availability of appropriate tools for sophisticated, high-throughput genetic engineering. This is particularly problematic for microbes that will deployed in environments where antibiotic selection is undesirable and impractical (industrial bioproduction and agricultural applications). Most common methods used to transfer heterologous DNA into PGPR (*e.g.* replicating plasmids, homologous recombination-based allelic exchange, transposon mutagenesis) have substantial caveats that limit their utility for engineered microbes in moderate- to high-throughput environmental and plant-microbiome interaction studies. Most replicating plasmids, including commonly used broad-host variants (*e.g.* pBBR^11,12^, RK2^13,14^, RSF1010 ^15^, and pBC1^16^ plasmids) are unstable ^17,18^ and incur a fitness cost ^19,20^ – thus requiring constant selection (typically antibiotic resistance) for maintenance. Furthermore, plasmid copy number variation both within a species ^21-23^ and between species ^24,25^ adds uncertainty when interpreting results. Both Tn and homologous recombination-based technologies stably integrate heterologous DNA into the genome ^26^. Despite high DNA integration efficiency, the unpredictable location of transposition events precludes simple comparison between pathway variants. Homologous recombination-based allelic exchange relies upon endogenous DNA-repair machinery – which is typically inefficient ^27^, precluding the construction of even moderately-sized strain libraries.

Integrase-mediated recombination technologies ^28-37^ have the potential overcome many of the limitations faced by the above genetic tools. Phage integrases (or site-specific recombinases) are enzymes that catalyze recombination between two specific sequences of DNA ^34,38^. Two tyrosine recombinases, Flp and Cre, are commonly used in molecular genetics^37,39^, but these enzymes recognize identical sites and perform reversible recombination. The other class of phage integrases, serine integrases, are single subunit enzymes that catalyze unidirectional DNA recombination between two short, distinct *att* sites (*attP* and *attB*), generating new attachment sites (*attL* and *attR*) in the process. Furthermore, serine integrases, such as Bxb1 and ΦC31, have been used across the tree of life and thus are likely to work in the vast majority of organisms. Phage integrase-based tools are often limited to integration of a single DNA fragment ^28,29,31,32,35,36,39^ or function in a limited range of hosts ^30,33,40^. To address these issues and limitations, in this work we developed the <u>S</u>erine-integrase <u>A</u>ssisted <u>G</u>enome <u>E</u>ngineering (SAGE) system. SAGE utilizes 10 serine integrases to enable efficient, iterative integration of 10 distinct DNA fragments into bacterial genomes at 10 unique *attB* sites (**Fig. 1**). We demonstrate robust SAGE performance in two phylogenetically distant PGPR, *Pseudomonas fluorescens* SBW25 and *Rhodopseudomonas palustris* CGA009. Furthermore, we highlight the utility of SAGE by developing a constitutive, genome-integrated promoter library that enables tunable (up to ∼40,000-fold dynamic range) expression of heterologous proteins in *P. fluorescens*.

**Fig. 1.**
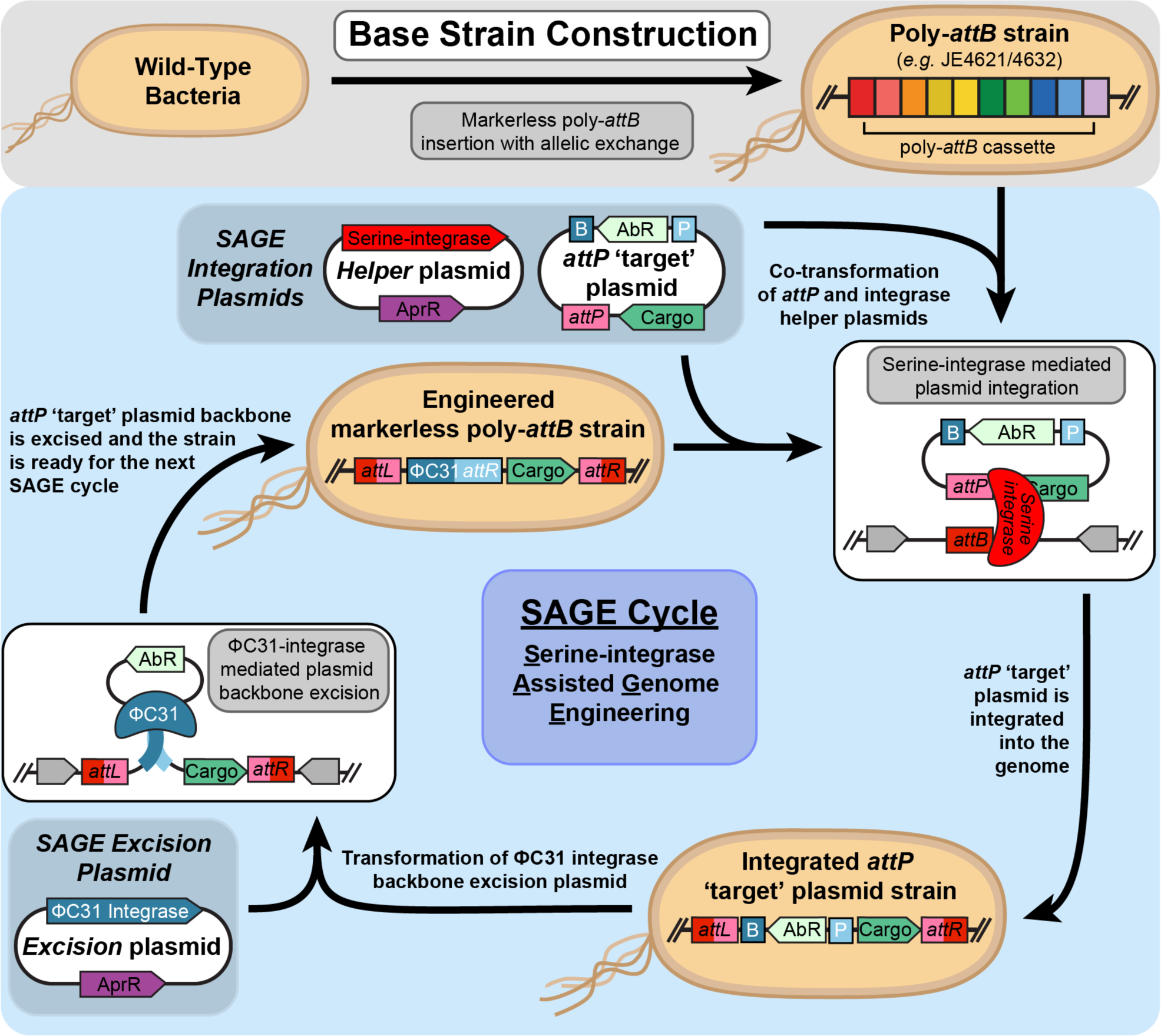
Iterative integration of multiple genetic constructs into the chromosome using <u>S</u>erine-integrase <u>A</u>ssisted <u>G</u>enome <u>E</u>ngineering (SAGE). Schematic performing cycles of SAGE, a technology for rapid, high efficiency, unidirectional, and site-specific integration of multiple DNA fragments into the chromosome of microbes. In the first step, base strain construction, *attB* sequences (in this case a 10-member poly-*attB* cassette) are integrated into the target microbe’s genome using standard methods (*e.g.* allelic exchange). Next, one of the system’s 10 integrases is transiently expressed by from a non-replicating *helper* plasmid, and catalyzes recombination between its cognate *attP* (co-transformed ‘target’ plasmid) and *attB* (chromosome) sequences. This unidirectionally integrates the plasmid into the genome and generates new *attL* and *attR* sequences. Next, ΦC31 integrase is transiently expressed by introducing a conditionally replicating or non-replicating *excision* plasmid and catalyzes recombination between ΦC31 *attP* (P) and *attB* (B) sites on the integrated ‘target’ plasmid. This excises both the selection marker and *Echerichia coli* origin of replication from the chromosome while leaving the integrated DNA cargo in the genome. As the excised circular DNA cannot replicate in organisms that are not closely related to *E. coli* the resulting strain is ready to be further engineered by using one of the SAGE system’s other 9 serine integrases.

## Results

### Development of a conditional-replication plasmid for transient protein expression in soil Pseudomonads

Many *Pseudomonads* are PGPR and have been used in commercial fertilizer products (e.g. *P. fluorescens*^41,42^ and *P. protegens*^43,44^). Development of high-throughput genetic tools to engineer additional functionality or improve performance of *Pseudomonad* and other PGPR isolates has substantially lagged behind those of related strains used for industrial fuels and chemicals production (e.g. *P. putida*^*45,46*^).

Conditional-replication plasmids are valuable for genetic engineering and investigation of essential genes. For example, plasmids with temperature-sensitive origins of replication (*ts* plasmids) deliver recombineering machinery and CRISPR/Cas systems for genome engineering in *Escherichia coli*^47,48^ and can be used for conditional knockout assays for essential genes. We developed a *ts* plasmid, pGW26, for plant growth-promoting (PGP) *Pseudomonads*, and demonstrate its utility for transient expression of serine integrases, henceforward referred to as integrases. pGW26 was constructed using a temperature-sensitive *Pseudomonas* origin of replication (mSF^*ts*1^)^49^, a pUC origin for replication in *Escherichia coli*, and an antibiotic resistance marker (**Fig. 2a**). The original mSF^*ts*1^ plasmid utilized ampicillin-selection, but as most *Pseudomonads* are natively resistant to ampicillin we utilized an apramycin resistance marker^27^.

**Fig. 2.**
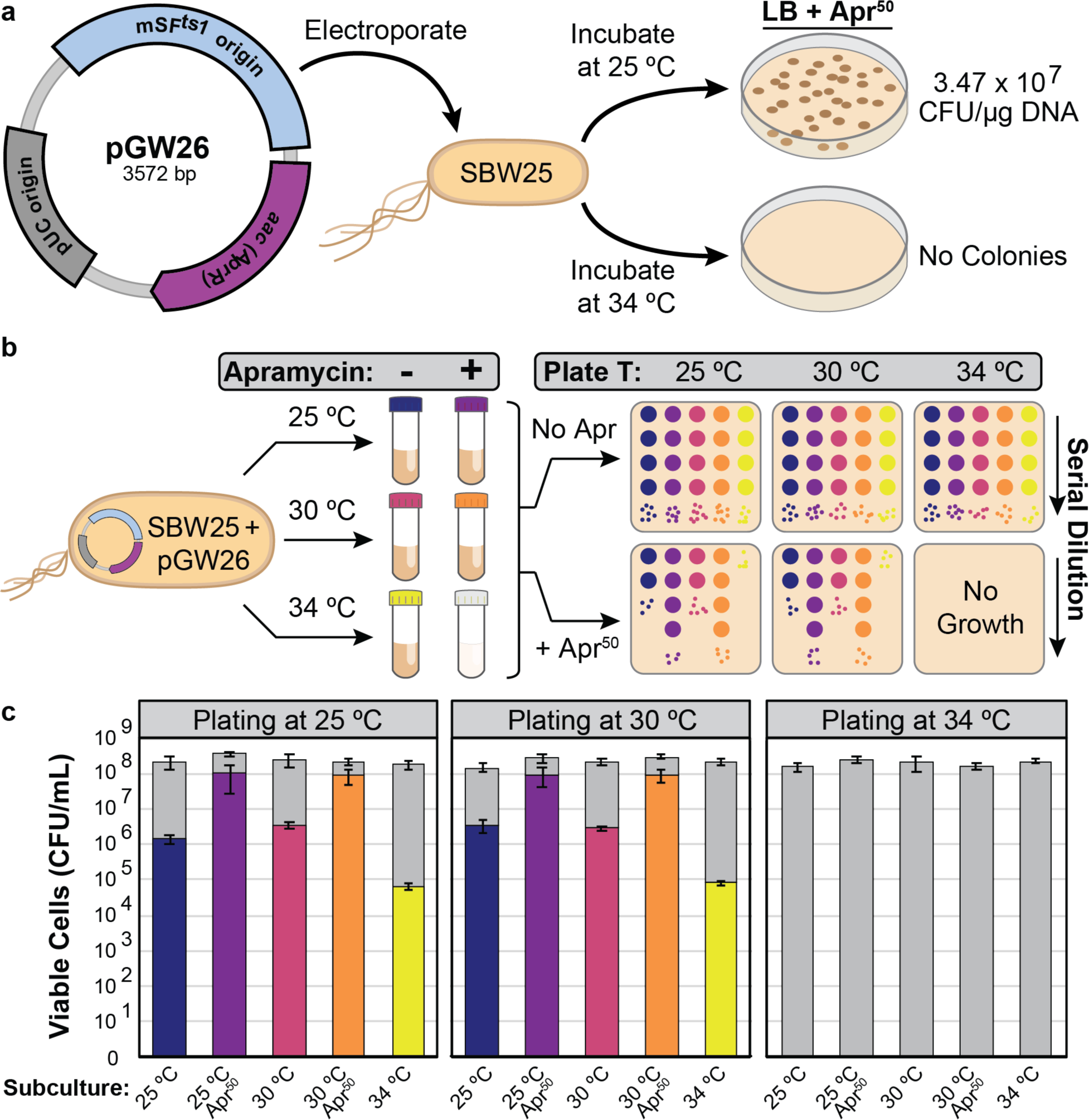
Development of a conditionally-replicating plasmid for broad use in *Pseudomonads*. (**a**) Graphic diagram of the temperature-sensitive pGW26 plasmid and an initial assay for determining its sensitivity to replication at 34 °C in *Pseudomonas fluorescens* SBW25. (**b**) Graphic diagram of a plasmid stability assay to determine the stability of pGW26 under growth without selection at temperatures that are ‘permissive’ and ‘non-permissive’ for plasmid replication. SBW25 containing pGW26 was cultivated with apramycin selection at 25 °C, and subsequently subcultured in selective or non-selective liquid medium at three different temperatures. Each resulting culture was evaluated for total culture viability (cell viability with no apramycin) or plasmid maintenance (cell viability with apramycin) at each of the three temperatures. No growth was observed in subculture in selective medium at 34 °C so it is not included in the subsequent plasmid maintenance assay. (**c**) Charts displaying results of plasmid stability assay. Gray bars indicate viable cells per mL in each culture, and colored bars indicate viable cells with apramycin selection (*i.e.* cells hosting plasmid). Error bars represent standard deviation in four replicate cell viability measurements.

Silo-Suh, *et al* originally identified 25 °C and 37 °C as permissive and non-permissive temperatures for mSF^ts1^ origin replication^49^, but intermediate cultivation temperatures were not analyzed. Many environmental *Pseudomonads*, including SBW25, cannot growth above 34 °C ^50,51^. So, we tested whether pGW26 would replicate and confer apramycin resistance to SBW25 at either 25 or 34 °C. When cultivated at 25 °C transformation was robust, (**Fig. 2a**) with colonies forming by 24 hours. However, as indicated by no growth after 72 hours, the plasmid likely does not replicate at 34 °C.

We next assessed plasmid stability by incubating a strain hosting pGW26 at three temperatures (25, 30, and 34 °C) in the presence or absence of antibiotic selection (**Fig. 2b-c**). If the plasmid is somewhat stable growth should occur in the presence of apramycin. Transformed SBW25 grew under all conditions except at 34 °C with antibiotic selection. We assessed cell viability (no apramycin) and plasmid maintenance (with apramycin) in each of the other five cultures by incubating serial dilutions of each on solid medium at 25, 30, and 34 °C.

Unsurprisingly, cell viability was similar for all five subcultures (**Fig. 2c, gray bars**), but we observed substantial differences in plasmid maintenance (**Fig. 2c, color bars**). Plasmid was retained in 50% of cells in the “permissive” temperature subcultures (25 and 30 °C) that contained antibiotic. The plasmid-bearing strain lost the apparently highly unstable plasmid at high-rates in all cultures lacking antibiotic. However, plasmid loss was 37-fold higher at 34 °C (99.96%) than at “permissive” temperatures (98.4-98.6%). Regardless, the absence of colonies on selective medium incubated at 34 °C demonstrates that pGW26 is readily cured in a single passage.

### Multiple serine integrases enable highly efficient unidirectional genomic integration of DNA

We next assessed the performance of <u>S</u>erine-integrase <u>A</u>ssisted <u>G</u>enome <u>E</u>ngineering (SAGE) with *P. fluorescens* SBW25 derivatives containing a genome integrated poly*-attB* cassette (**Fig. 3**). The cassette contains distinct *attB* sequences for 10 integrases (**Fig. 3a**), and was integrated into two putative “safe sites” in the genome via allelic exchange ^52^. The cassette was integrated downstream of either *ampC*, a class C β-lactamase, (strain JE4621) or PFLU5798, a TonB-dependent siderophore receptor (strain JE4624).

**Fig. 3.**
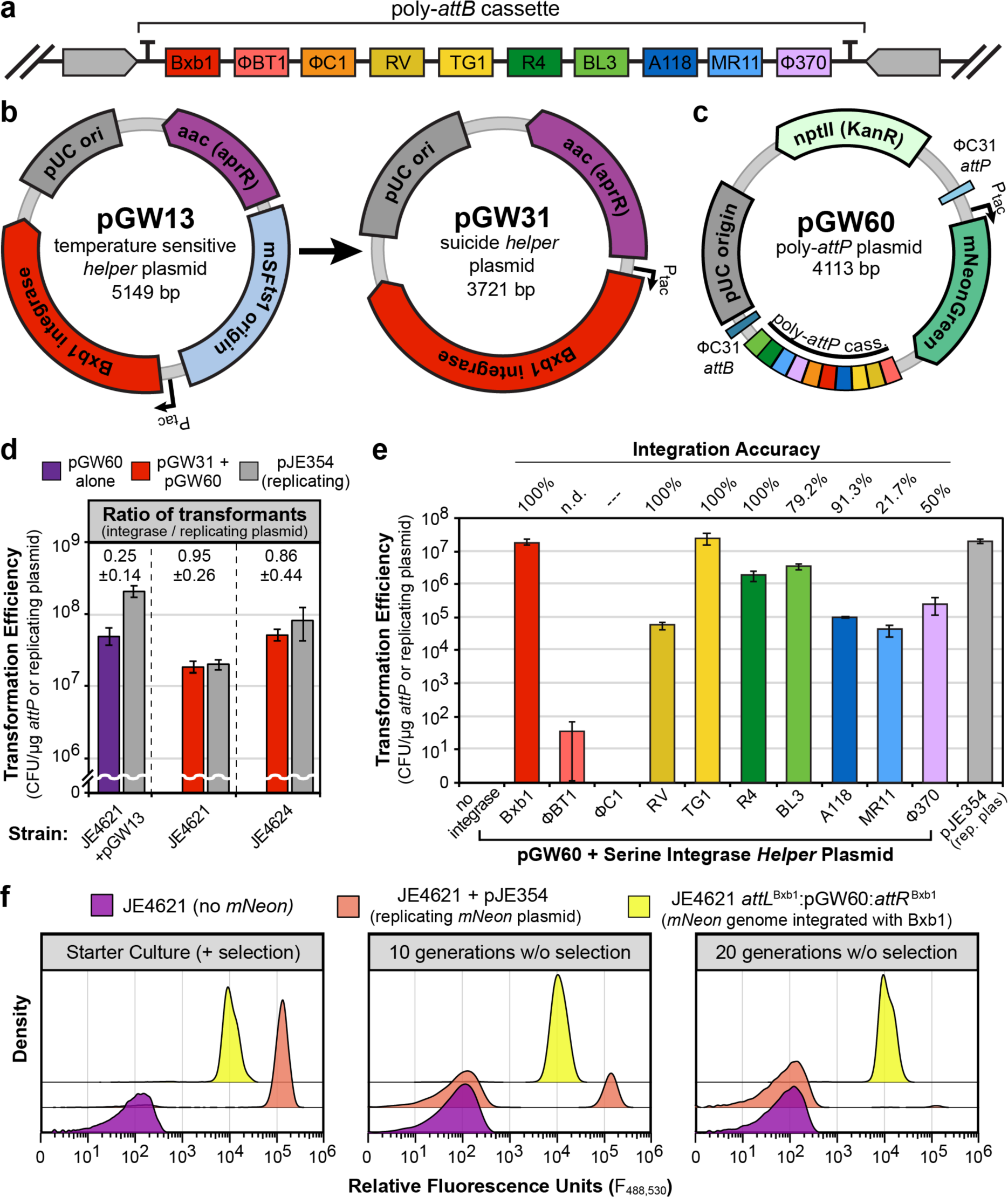
Multiple serine-integrases enable stable, highly efficient integration of plasmid DNA into the genome of engineered *P. fluorescens*. (**a**) Diagram of genome-integrated 10x poly-*attB* cassette, which includes flanking double terminators for transcriptional insulation. Each *attB* sequence is indicated by a color-coded box and internal label of associated integrase, and is flanked by a random DNA spacer sequence. Plasmid maps of Bxb1 integrase delivery plasmids (**b**) and poly-*attP* ‘target’ plasmid (**c**). Colored boxes in poly-*attP* cassette indicate the associated integrase for each *attP* in the array. (**d**) Transformation efficiency with a replicating plasmid (gray) or integrase target plasmid where Bxb1-integrase is delivered by a previously-transformed *ts* plasmid (purple) or co-transformed *suicide* vector (red). (**e**) Transformation efficiency using replicating plasmid pJE354, pGW60 alone, or pGW60 co-transformed with each of the integrase suicide *helper* plasmids. Bars are color-coded by integrase. (**d-e**) Error bars indicate standard deviation in three replicate assays. (**f**) Flow cytometry analysis of the parent strain (purple), and strains expressing mNeonGreen from a replicating plasmid (orange) or genome integrated copy (yellow). Y-axis is the distribution of cells with a given relative fluorescence level in the population.

We initially assessed performance of Bxb1 integrase – the best performing integrase in *P. putida*^29^. A constitutive Bxb1-integrase expression cassette was inserted into pGW26, generating pGW13 (**Fig. 3b**), and the resulting plasmid was transformed into JE4621. We assessed performance of Bxb1-integrase by electroporating this strain with either a pBBR1-MCS-2^11^ derivative, pJE354 (**Supplementary Fig. S1**), or pGW60, a non-replicating *suicide* plasmid that contains 10 *attP* sequences - one cognate *attP* for each of the genome integrated *attB* sequences (**Fig. 3c**). Both plasmids harbor constitutive mNeonGreen expression cassettes to visually assess transformation. Transformants with pJE354 are expected to contain autonomously replicating plasmid. Transformants with pGW60 are expected to result from unidirectional recombination between Bxb1-*attP* (pGW60) and Bxb1-*attB* (genome), thereby generating two new *att* sequences, Bxb1-*attL* and Bxb1-*attR* (**Fig. 1**).

Transformation efficiency was measured by enumerating kanamycin resistant colonies (**Fig. 3d**). Transformation efficiency with the replicating plasmid pJE354 was very high (2.07 × 10^8^ CFU/µg DNA). We observed similar, though slightly lower transformation (∼25%) efficiency with pGW60, suggesting that Bxb1 integrase expressed from pGW13 enables highly efficient integration of Bxb1 *attP*-containing plasmid DNA into the genome. Of note, no colonies were produced by electroporation of the Bxb1 integrase-null strain, JE4621, with pGW60 alone.

In principle, expression of the integrase should only be required during the recovery cultivation following electroporation, and thus the requirement for using a *ts helper* plasmid to express the integrase in the initial SAGE protocol may be dispensable (**Supplementary Fig. 2a**). Specifically, we hypothesized that simultaneous electroporation of pGW60 with a second non-replicating (*suicide*) integrase expression *helper* plasmid would result in efficient integration of pGW60 into the genome. This would reduce SAGE cycle time by at least 2 days (**Supplementary Fig. 2b**), and will help with adapting SAGE to organisms lacking easily curable plasmids or few antibiotic resistance markers.

We tested this hypothesis by removing the *Pseudomonas* origin or replication (mSF^ts1^) from pGW13, generating pGW31, and attempting transformation of the poly-*attB* strain JE4621 by simultaneously electroporating both pGW31 and pGW60 (**Fig. 3d**). Of note, batch-to-batch variance in competence can range over an order of magnitude, so the ratio of replicating plasmid to integrated pGW60 transformants within a given strain can be more informative than raw transformant counts. This is particularly important when comparing results between strains. Indeed, when Bxb1 integrase was delivered by a non-replicating *helper* integration of pGW60 was on par transformation efficiency of pJE354. We also observed similar results in an identical experiment with JE4624 as the recipient strain (**Fig. 3d**), demonstrating that *attB* location has minimal impact on integrase performance. This clearly demonstrates that transient expression of Bxb1 integrase is sufficient for high-efficiency DNA integration.

While raw transformation values varied by strain, and is likely an artifact of variable quality for each competent cell preparation, the ratio of pGW60 to pJE354 transformants was substantially lower (0.25 ± 0.14) when Bxb1 integrase was delivered by the replicating plasmid than when delivered via co-transformation. This suggests two things: 1) the previously observed loss of *ts* plasmid during growth with apramycin selection reduces the number of cells with sufficient Bxb1 expression for DNA expression, and 2) uptake of two distinct plasmids is as efficient as uptake of a single plasmid in electrocompetent *P. fluorescens* SBW25 cells.

A single efficient integrase can enable construction of strain libraries with up to 5×10^7^ members (**Fig. 3d**), but a given integrase can only be utilized once in a strain. Use of multiple integrases would enable production of large combinatorial strain libraries, and many serine integrases have been described in the literature (see reviews^34,38,39^). We hypothesized that additional serine integrases would function well in *P. fluorescens*, and tested nine more with the integrase expression *helper* plasmid / pGW60 co-transformation method described above (**Fig. 3e**). With two exceptions (BT1 and ΦC1), most additional serine integrases were able to efficiently catalyze recombination between their respective *attP* (in pGW60) and *attB* (genome) sequences, resulting in high transformation efficiencies (4.41 × 10^4^ to 2.53 ×x 10^7^ CFU/µg of pGW60 DNA).

Integration accuracy, an important metric when assessing the utility of each integrase as a genetic tool, is the frequency of recombination at the intended *attB* site versus off-target integration at ‘pseudo-*att*’ sites. The characterized *attP* and *attB* sequences are derived from the phage recombination sites in the originally characterized bacteriophage-host microbe combination, but may not represent the only, or even the highest affinity, site for integrase-mediated recombination in a given organism. Accordingly, the presence of pseudo-*att* sites is dependent on chromosome sequence, and needs to be assessed in each new organism. We performed a colony PCR assay using both a pGW60 integration-specific primer set and a control primer set to assess integration accuracy. Given their poor performance in *P. fluorescens* we did not test ΦC1 or BT1 in this or following assays. Four of the eight remaining integrases, including Bxb1 and TG1, displayed 100% (24/24) accuracy (**Fig. 3e**). A118 and BL3 were relatively accurate, but MR11 and Φ370 did not integrate at the target locus more than 50% of the time – making them suboptimal for SAGE in *P. fluorescens*.

### Genomic integration of heterologous DNA with SAGE enables stable gene expression

A critical advantage SAGE has over replicating plasmids is consistent, stable expression in the absence of antibiotic selection. This is particularly important for engineering PGPR that will be deployed in environments where standard selections (*e.g.* antibiotics, auxotrophies) are non-viable. We hypothesize that engineered functions will be more stable when introduced with the integrase system versus a replicating plasmid.

As a proxy for engineered functions we assessed the stability of fluorescent protein expression. *P. fluorescens* JE4621 derivatives hosting mNeonGreen expression cassettes on either a replicating plasmid, or integrated into the genome at each of the *attB* sites (with the exception of BT1 or ΦC1) were assayed for stability mNeonGreen production in the absence of antibiotic selection using flow cytometry. Irrespective of integrase utilized for integration, expression of mNeonGreen was stable in all strains where pGW60 was integrated into the genome (**Fig. 3f – yellow, Supplementary Fig. S3**). Despite cultivation in presence of antibiotic, 2.5% of cells in the replicating plasmid starter culture lacked mNeonGreen expression. After 20 generations in the absence of antibiotic, 97.5% of cells failed to produce mNeonGreen (**Fig. 3f – orange**) This demonstrates the unstable nature of plasmid-based engineered functions. Furthermore, the inconsistency of mNeonGreen expression in an antibiotic-containing starter culture suggests that some plasmid loss occurs during cultivation with antibiotic selection.

### *Transient expression of* ΦC31 *integrase enables selection marker recycling and iterative genome engineering*

The ability to excise, and thus recycle, selection markers with a second phage integrase enables their use for iterative rounds of SAGE (**Fig. 4**). The ΦC31 integrase is a widely used integrase {Brown, 2011 #123;Guss, 2008 #116;Keravala, 2006 #117;Xu, 2016 #119}, and thus is likely to function in most organisms. Accordingly, the ‘cargo region’ of the pGW60 plasmid – containing mNeonGreen – is flanked by a ΦC31 *attP*/*attB* pair (**Figs. 3c, 4a**), and we hypothesized that if ΦC31 is functional in *P. flourescens* transient expression of the integrase should excise the selection marker and remainder of the ‘plasmid backbone’ (**Fig. 4b**). We tested this by transforming a strain where pGW60 was integrated into the genome, JE4689 (JE4621 *attL*^Bxb1^:pGW60:*attR*^Bxb1^, **Fig. 4a**), with a *ts* plasmid either contains (pGW30) or lacks (pGW26) a ΦC31 integrase expression cassette. Transformants were cured of their respective plasmids by cultivation at 34 °C on non-selective solid medium and screened for both the loss plasmid (colony PCR, apramycin sensitivity) and excision of the plasmid backbone (colony PCR). Colony PCR (**Fig. 4c**) and antibiotic resistance confirmed *ts* plasmid loss. However, the pGW60 plasmid backbone was only excised in pGW30 transformants (**Fig. 4d**), demonstrating that the selection marker can be recycled via ΦC31 excision.

**Fig. 4.**
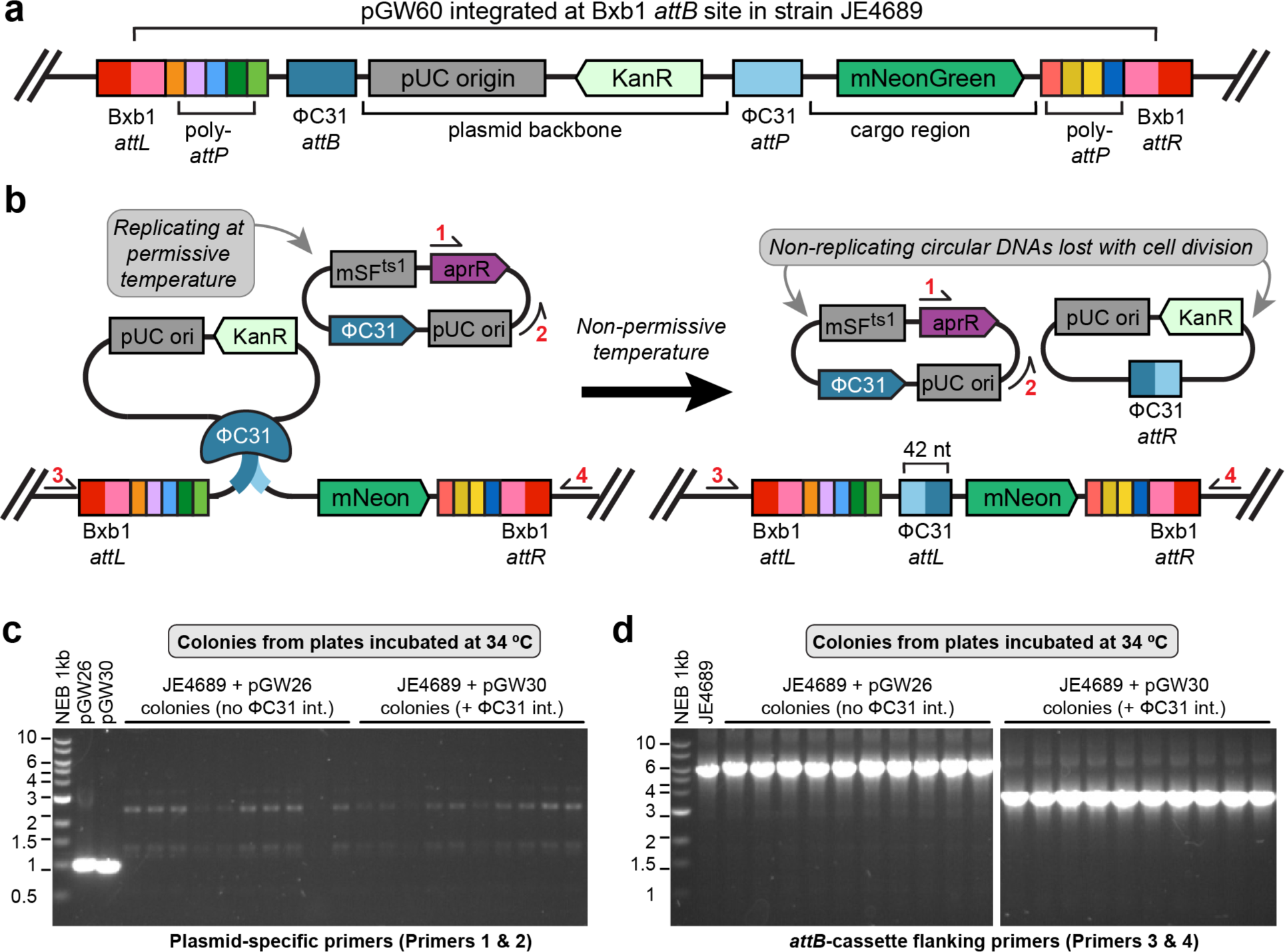
ΦC31-integrase mediated excision of ‘target’ plasmid backbones enables selection marker recycling. (**a**) Diagram of the genome of JE4621 with pGW60 integrated at the Bxb1-*attB* site. (**b**) Diagram of pGW60 plasmid backbone excision using a *ts helper* plasmid to deliver ΦC31-integrase. Colony PCR primers are indicated by arrows and red numbers. (**c-d**) Colony PCR validation of *ts* plasmid loss (**c**) and pGW60 plasmid backbone excision (**d**) following overnight cultivation of apramycin-resistant colonies at 34 °C.

### SAGE enables high-throughput analysis of genome integrated promoters

Expression of engineered pathways (N-fixation, secondary metabolite production, *etc*.) must be balanced with core cellular functions to minimize impact on host fitness and increase their evolutionary stability. Soil conditions vary substantially over time and location, so it is important to identify a set of promoters for each PGPR that covers a wide range of expression levels that perform consistently across diverse conditions. Promoter libraries have been developed using reporter protein expression in several organisms ^53-56^. However, most libraries, including the commonly used ‘Anderson’ library (http://parts.igem.org/Promoters/Catalog/Anderson), were developed using multicopy reporter plasmids. Heterogeneity in copy number among colonies, within culture subpopulations (**Fig. 3f**), as well as overall higher DNA dosage can influence reported relative promoter performance. When reporter constructs are genome-integrated the copy number is lower and more consistent. Furthermore, the host is free of any metabolic burden imposed by plasmid maintenance. So, plasmid-based promoter performance measurements may not always correlate well with those generated using genome-integrated reporters.

Accordingly, we developed a method to rapidly characterize large libraries of genome-integrated promoters that combines the efficiency of SAGE integration with a modified version of the high-throughput sequencing (HTS) transcription profiling method described by *Yim, et al* ^57^. We apply this method, described briefly below, to characterize a collection of 287 synthetic and natural promoters (**Supplementary File T1**) in *P. fluorescens* (**Fig. 5a**). Of note, we modified the original method in two important ways. First, rather than a single barcode per promoter, we generated 5 distinct barcode variants per promoter. Second, the barcode was located upstream of the RBS in the 5’ UTR, rather than in the reporter coding sequence. This enables analysis of promoter (in)sensitivity to 5’ UTR perturbation – an important metric for heterologous promoters, as 5’ UTR sequence affects mRNA stability and it is unlikely that the same 5’UTR will always be utilized for both promoter characterization and promoter application. Second, with multiple barcodes we can have higher confidence in data for each promoter, and have the potential to identify and discard spurious outlier barcode variants.

**Fig 5.**
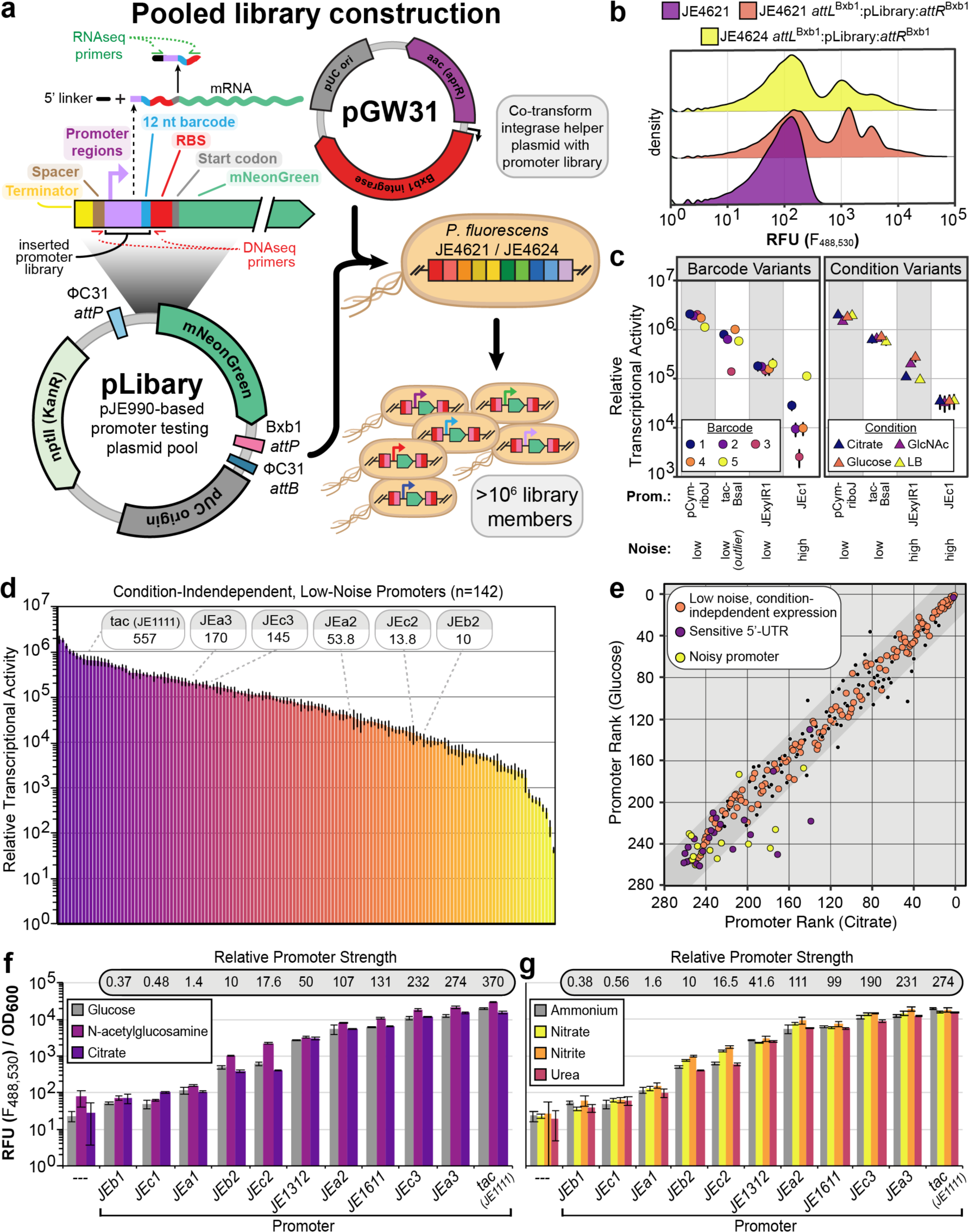
SAGE enables high-throughput construction and assessment of pooled genome integrated promoter libraries. (a) Overview of promoter library construction and DNA/RNAseq barcode sequencing fragments. (b) Flow cytometry evaluation of promoter library distribution in two *P. fluorescens* backgrounds. (c-e) Data from high-throughput sequencing promoter analysis assays. (c) Example promoters with sensitive 5’ UTR (high barcode-associated ‘noise’) and differential expression across conditions (condition-associated ‘noise’). (d) Chart displaying mean relative promoter strength – all barcodes and conditions averaged – for condition-independent, low noise promoters. Relative strength of the promoters used in panels f,g are indicated. (c-d) Error bars represent standard error in (c) 12-16 samples or (d) 40-80 samples. (e) Chart comparing ranking of RTA for each promoter during growth with glucose (y-axis) or citrate (x-axis) as sole carbon source. Dark gray band represents the range of values in which promoters must lie to be considered condition-independent for each pair-wise condition comparison. Colors indicate the classification of each promoter. (f-g) Promoter performance for a small subset of pLibrary promoters and *P. putida* KT2440 promoter library members (JE1312, JE1611). Error bars represent the standard deviation in 3 replicates. Relative promoter strength is calculated by comparing mean RFU/OD600 values across all (f) carbon or (g) nitrogen sources. (g) Glucose and (f) ammonium values in are from the same samples.

SAGE was utilized to integrate the pooled library of barcoded reporter plasmids (pLibrary) into the genomes of both *P. fluorescens* JE4621 and JE4624, generating two pooled libraries of reporter strains, with each transformant containing an integrated pLibrary plasmid. The JE4621 parent strain (no library), JE4621 and JE4624 libraries, as well as the *E. coli* plasmid host strain library were analyzed by flow cytometry (**Fig. 5b** and **Supplementary Fig. 4**). The JE4621 and JE4624 libraries had similar fluorescence profiles when analyzed by flow cytometry, which suggests that the site of integration does not substantially impact library performance. Next, we cultivated the JE4621-based library in several different media conditions to identify which promoters are both insensitive to 5’-UTR perturbations and provide condition-independent expression – a essential trait for constitutive promoters to drive reliable expression of engineered pathways in heterogenous environments. Specifically, we cultivated the JE4621-based library in quadruplicate in either a rich medium (LB) or defined mineral medium (M9) supplemented with one of three soil-relevant carbon sources: glucose (cellulose), citrate (root exudate), or N-acetylglucosamine (chitin). Next, we performed targeted high-throughput sequencing to enumerate barcode abundance in the DNA and RNA fractions from each sample. Relative transcriptional activity in a given sample was calculated by comparing the RNA:DNA barcode abundance ratio.

Of the 287 promoters, 262 had at least three barcode variants present in all 16 samples and were further analyzed (**Supplementary Table T2**). We evaluated promoters for 5’ UTR sensitivity by comparing the relative transcriptional activity (RTA) between barcode variants of the same promoter (**Supplementary Fig. 5**). We found that 25 of the 262 promoters displayed high variation among the barcode variants (see **Fig. 5c**, promoters pCym-riboJ and JEc1 for examples of 5’-UTR insensitive and 5’UTR-sensitive promoters, respectively). In many cases a single barcode variant was an outlier from the others (see **Fig. 5c**, promoter tac-BsaI). These results highlight the importance of considering the impact of barcode sequences have upon expression when performing analyses of regulatory elements, and that the use of multiple barcodes will be critical to reliably assess performance in high-throughput assays. We identified an 15 additional promoters with noisy expression (**Supplementary Fig. 5**) when analyzing the RTA data with a condition and barcode agnostic method.

We next evaluated the promoters for consistent expression between conditions (**Supplementary Figs. S6 and S7**). We ranked each promoter from highest (rank 1) to lowest (rank 262) RTA in each condition performed pairwise comparisons between each condition (see **Fig. 5e** and **Supplementary Fig. S6**). Promoters whose rank shifted by less than 10% of the library size (26 positions) for each pair of conditions was considered to have condition-independent expression (**Fig. 5e** – orange dots). Using this metric 142 of the 222 low noise promoters were considered condition-independent.

RTA values of the low noise, condition-independent promoter collection ranges over a 43,080-fold range of expression (**Fig. 5d**), with 90% within a 716-fold range of expression. Interestingly, there was a general trend of the weakest promoters displaying a somewhat bimodal pattern of expression between samples – with promoters either expressed at moderately low expression levels or with no expression in a given sample (**Supplementary Fig. 5b**). This bi-modality may be a consequence of transcriptional bursting ^58^ switching a promoter from *on* to fully *off* in a stochastic manner, and if so would suggest that weaker promoters may be inherently noisier as a consequence of the phenomenon.

We validated the general trends in relative promoter strength from the HTS-based assay with microtiter plate fluorescent reporter assays. Reporter plasmids containing representative promoters from this library and a *Pseudomonas putida* genome-integrated promoter library (P_JE1312_ and P_JE1611_) ^29^ were integrated into JE4621 using SAGE. The resulting strains were assayed for mNeonGreen production in M9 medium containing ammonium with three distinct carbon sources (glucose, citrate, N-acetylglucoseamine) or glucose with four distinct nitrogen sources (ammonium, nitrate, nitrite, urea) in a microtiter plate growth assay. Relative promoter strength for was consistent across all 7 tested conditions (**Fig. 5f**). With the exception of the three weakest promoters (P_JEa1_, P_JEb1_, P_JEc1_), which were noisy promoters in the library data, the relative promoter strengths for the other promoters were very similar to those from the HTS-based assay (**Figs. 5d**,**f**). Furthermore, when normalized to JE1111 (tac) the relative performance of P_JE1611_ (0.36 vs 0.3) and P_JE1312_ (0.14 vs 0.12) are exceptionally similar to their performance in *P. putida* KT2440 – suggesting that our promoter library may translate well to the industrially-relevant microbe.

### SAGE enables efficient genome engineering in diverse bacteria

Serine-integrases are functional in a wide array of eukaryotes, archaea, and bacteria^28-33,35,36,40,59^ – so SAGE should work in any transformable bacterium where two conditions are met: *att* sites are integrated into the genome (**Fig. 1**) and active serine integrases are expressed. We assayed SAGE in the phylogenetically distant alphaproteobacterium, *Rhodopseudomonas palustris*^*60*^. Able to grow under both anaerobic and aerobic conditions in heterotrophic, phototrophic, and chemolithotrophic modes of growth, *R. palustris* has one of the most versatile metabolic repertoires known bacteria, and are also often PGPR^61-64^.

We replacing RPAL1300, which encodes a Type IV restriction endonuclease, with the poly-*attB* cassette to generate a derivative of *R. palustris* CGA009^*60*^, JE4632, that we could to assess SAGE performance. Each integrase was assayed in JE4632 using plasmid co-transformation assays as described above for *P. fluorescens* JE4621 (**Fig. 3e**). In addition to a replicating plasmid control, pEYF2K, we also included an additional control, pJE1609. Transformants using pJE1609 arise from recombination catalyzed by the endogenous DNA repair machinery at one of two 700 bp regions of homology between the plasmid and genome.

All 10 integrases were efficient at integrating pGW60 into the JE4632 genome (**Fig. 6a**), but we noticed some notable differences in integrase performance compared to *P. fluorescens* JE4621 (**Fig. 3e, Supplementary Fig. S8**). Transformation efficiency increased with 8 of 10 integrases, and increased by >10^2^ with five integrases (ΦBT1, ΦC1, RV, A118, and R4). While these gains may be due in part to an overall increase in transformation efficiency, we observed a significant decrease in efficiency with two integrases (TG1 and BL3). Overall integrase accuracy was higher in JE4632, with MR11, BL3, and Φ370 improving substantially. However, TG1 integrase accuracy decreased to 62.5%, and may explain its decreased transformation efficiency in JE4632. Unsurprisingly, given the typical low efficiency of endogenous homologous recombination, pJE1609 transformation efficiency was orders of magnitude lower than any integrase. Finally, the poor performance of pEYF2K in comparison to the integrases was striking, and may be indicative of an insufficiently expressed selection marker or instability.

**Fig. 6.**
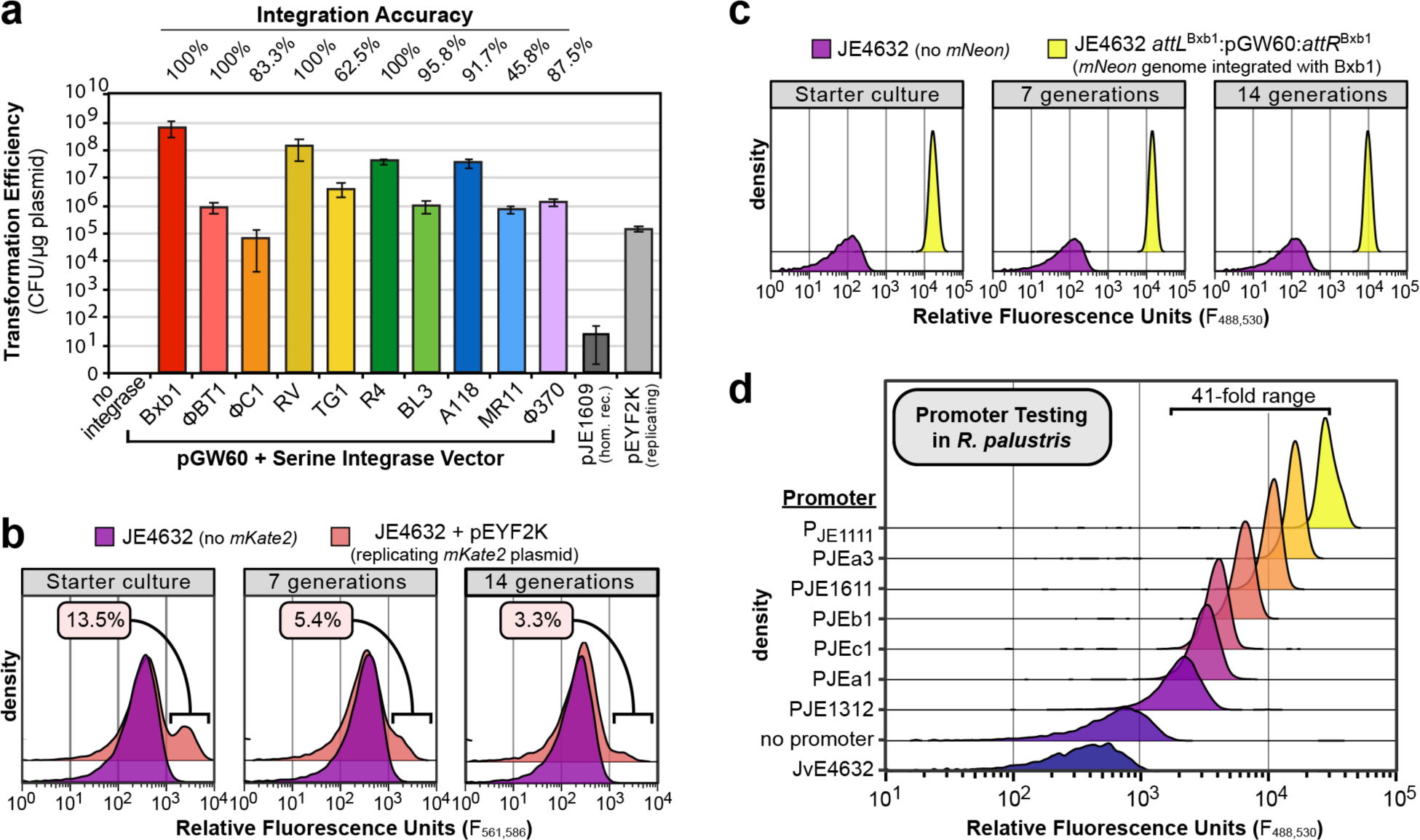
SAGE performance, plasmid instability, and a small promoter library in *Rhodopseudomonas palustris* CGA009. (**a**) Transformation efficiency using replicating plasmid pEYF2K, suicide homologous recombination plasmid pJE1609, pGW60 alone, or pGW60 co-transformed with each of the integrase suicide *helper* plasmids. Bars are color-coded by integrase or alternate plasmid classes (gray shades). Error bars indicate standard deviation in three replicate assays. (b) Flow cytometry analysis of the parent strain (purple) and strains expressing mKate2 from a replicating plasmid (orange). (c) Flow cytometry analysis of the parent strain (purple) and genome integrated copy (yellow). (d) Flow cytometry-based analysis of reporter protein expression using a small library of genome-integrated promoter testing constructs.

We next compared stability of heterologous DNA in the absence of selection with an experimental design similar to that used to test *P. fluorescens* JE4621 derivatives (**Fig. 3**), but with two minor alterations. First, we were unable to transform JE4632 with pJE354, potentially due to poor activity of the *aphI* kanamycin-resistance marker. Instead, we used pEYF2K, another pBBR1-plasmid with the *npt-II* kanamycin-resistance marker, and tracked its stability by monitoring expression of the red fluorescent protein mKate2. Second, as a consequence of its much slower growth, for *R. palustris* we performed flow cytometry analysis every 7 generations (**Fig. 6b-c, Supplementary Fig. S9**). Interestingly, despite cultivation of an isolated pEYF2K transformant in selective medium, mKate2 is only expressed in 13.5% of the starter culture cells (**Fig. 6b**). Nonetheless, further loss of fluorescence (and likely plasmid) is observed, and after 14 generations without selection mKate2 expression is further reduced to 3.3% of the population. However, expression of mNeonGreen is maintained in all cells where pGW60 was incorporated into the genome using SAGE (**Fig. 6c, Supplementary Fig. S9**).

We also utilized SAGE to assess performance of a small collection of genome integrated promoters. *R. palustris* is photosynthetic grows best when exposed to light under microaerobic or anaerobic conditions. Accordingly, we utilized flow cytometry analysis of microaerobic test tube cultures rather than microtiter plate cultures to assess mNeonGreen production in each reporter strain (**Fig. 6c**). Overall, the range of expression levels with this collection promoters was substantially narrower than in *P. fluorescens* (41-fold versus >726-fold). Interestingly, while the order of the three strongest promoters remains the same (P_tac_ > P_xylE_2_ > P_JE16111_), the relative strengths of the weaker promoters are substantially reordered relative to *P. fluorescens* – highlighting the need for a simple and efficient system to rapidly test genome integrated promoters in each new host.

### SAGE toolkit for scientific community

Using the knowledge gained here, we have developed a series of SAGE toolkit plasmids that are available for use (**Supplementary Fig. 10**). This toolkit includes both *suicide* (non-replicating) and mSF^ts1^ (temperature-sensitive *Pseudomonas* origin) integrase expression plasmids (pGW13-40 & pJE1817). It also includes a highly modular collection of single *attP* integrase ‘target’ plasmids that either contain the *nptII* (kanamycin resistance marker) alone or in combination with *sacB* for sucrose counter-selection. The *nptII*-*sacB* variants enable selection against maintenance of the plasmid backbone in the genome, and thus enable sucrose selection for plasmid backbone excision when using a *suicide* ΦC31 integrase *helper* plasmid. This removes the requirement to use a replicating plasmid for ΦC31 integrase delivery – thus speeding the backbone excision process (**Supplementary Fig. S2c-d**) and enabling excision in *sacB*-compatible bacteria for which replicating plasmids are unknown. All of the *attP* ‘target’ plasmids are modular and contain unique restriction sites flanking each major feature to enable easy swapping of selection markers, *E. coli* origins of replication, and *att* sites. The plasmids contain multiple cloning sites flanked by a pair of double rho-independent terminators that contain within multiple unique restriction sites – including a pair of BbsI restriction sites that are compatible with Golden Gate cloning. Plasmid pJE990, which we previously developed is a related plasmid that is setup as an easy to use out of the box reporter plasmid, with sites for integration of promoter elements, ribosomal binding site variants, and terminators – as well as convenient sites for replacing the reporter with any gene of interest. We expect that these tools, and future derivatives that address the specific needs for a given host range will enable SAGE to be applied throughout the bacterial domain.

## Discussion

SAGE is a powerful addition to the toolkit for high-throughput engineering of bacteria in any application space (*e.g.* nutraceuticals synthesis, biofuel production, *etc.*), and will be especially transformative for applications where selection for plasmid maintenance is impossible (*e.g.* soil or human gut microbiome engineering). The genome integration efficiency we observed when using SAGE in multiple phylogenetically distant bacteria is among the highest when compared to competing technologies, and has been used by our group to integrate >13 kb DNA fragments with no apparent loss in efficiency (data not shown). Furthermore, by utilizing a second serine-integrase (ΦC31) to excise the selection marker and plasmid backbone, SAGE enables iterative integration of up to 10 distinct pools of DNA fragments. Although not demonstrated here, serine-integrase mediated recombination is selectively reversible by co-expression of a serine-integrase with its cognate recombination directionality factor (RDF) protein ^65,66^. This reversibility could be harnessed for applications such as directed evolution where genome integrated plasmid variants could be “extracted” as circular DNA that can be transferred to *E. coli* for replication and further manipulation. Finally, SAGE is orthogonal to host recombination machinery and does not require use of any replicating plasmids, and thus is theoretically usable in any transformable bacteria – regardless of state of their current genetic tools. As long as a *attB* cassette can be transferred into the organism’s genome – via allelic exchange, recombineering, Tn-based integration, *etc*. – the bacteria should be amenable to SAGE.

## Supporting information

Supplementary File P1 - Plasmid Maps

Supplementary File T3 - Filtered HTS Transcription Data

Supplementary File T2 - Processed HTS Promoter Data

Supplementary File T1 - Promoter Metadata

## Acknowledgements

We would like to acknowledge Carrie Harwood and Yasuhiro Oda for providing *Rhodopseudomonas palustris* CGA009 and for helpful information regarding cultivation of the strain. We would also like to thank Enoch Yeung and Yuliya Farris for providing plasmid pEYF2K.

## Notes

### Competing Interest Statement

The authors have declared no competing interest.

