## Supplementary Figures for "The SAGE genetic toolkit enables highly efficient, iterative site-specific genome engineering in bacteria"

### Supplementary Figures and Tables

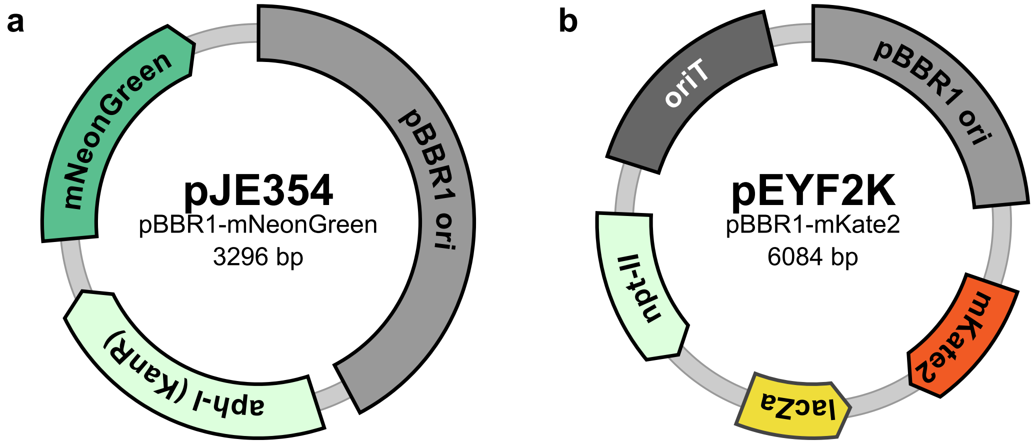


**Supplementary Fig. S1. Plasmid maps of replicating plasmids used in this study.** Both (a) pJE354 and (b) pEYF2K contain the widely utilized broad host-range pBBR1 origin that enables autonomous replication in many gram-negative bacterial hosts. (a) pJE354 is a derivative of pBTL-2 that contains a constitutive $P_{tac}$:mNeonGreen fluorescent protein reporter cassette, as well as the kanamycin resistance selection marker *aph-I*. (b) pEYF2K is a derivative of pBBR-MCS-2, contains a different kanamycin resistance marker than pJE354 (*npt-II* – of note, this is the same marker used in all KanR SAGE plasmids), a constitutive $P_{J23150}$:mKate2 fluorescent reporter cassette, an origin of transfer that enables conjugative transfer with the RK2/RP4 mobilization systems, and a lacZ$\alpha$ fragment for blue-white screening when cloning.

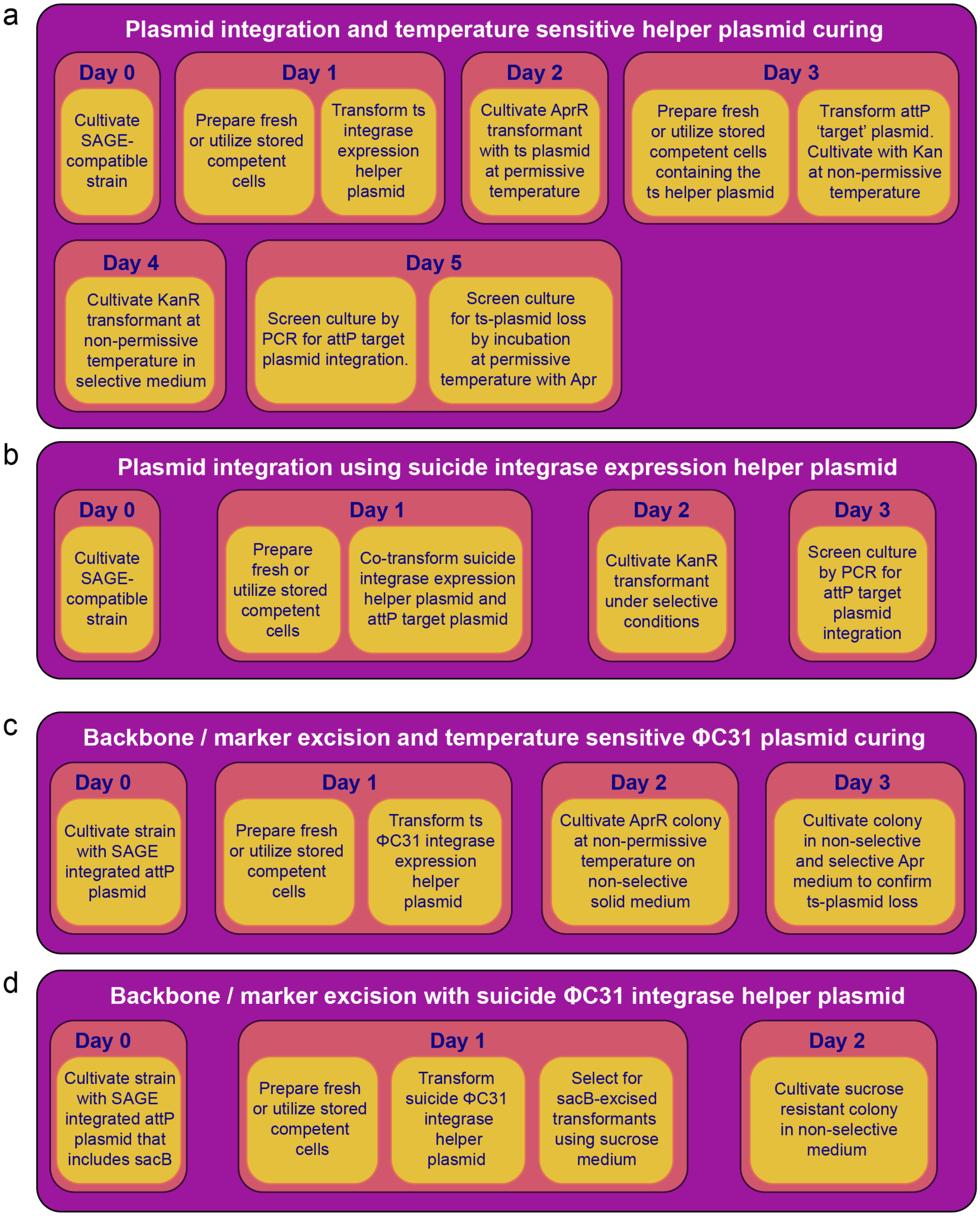


**Supplementary Fig. S2.** Overview timeline of steps when using different variants of SAGE for genome integration of *attP* target plasmids (a-b) and plasmid backbone excision with phiC31(c-d). (a,c) Protocol variants using temperature-sensitive *helper* plasmids to deliver the integrase expression cassettes. (b,d) Protocol variants using *suicide helper* plasmids to deliver the integrase expression cassette.

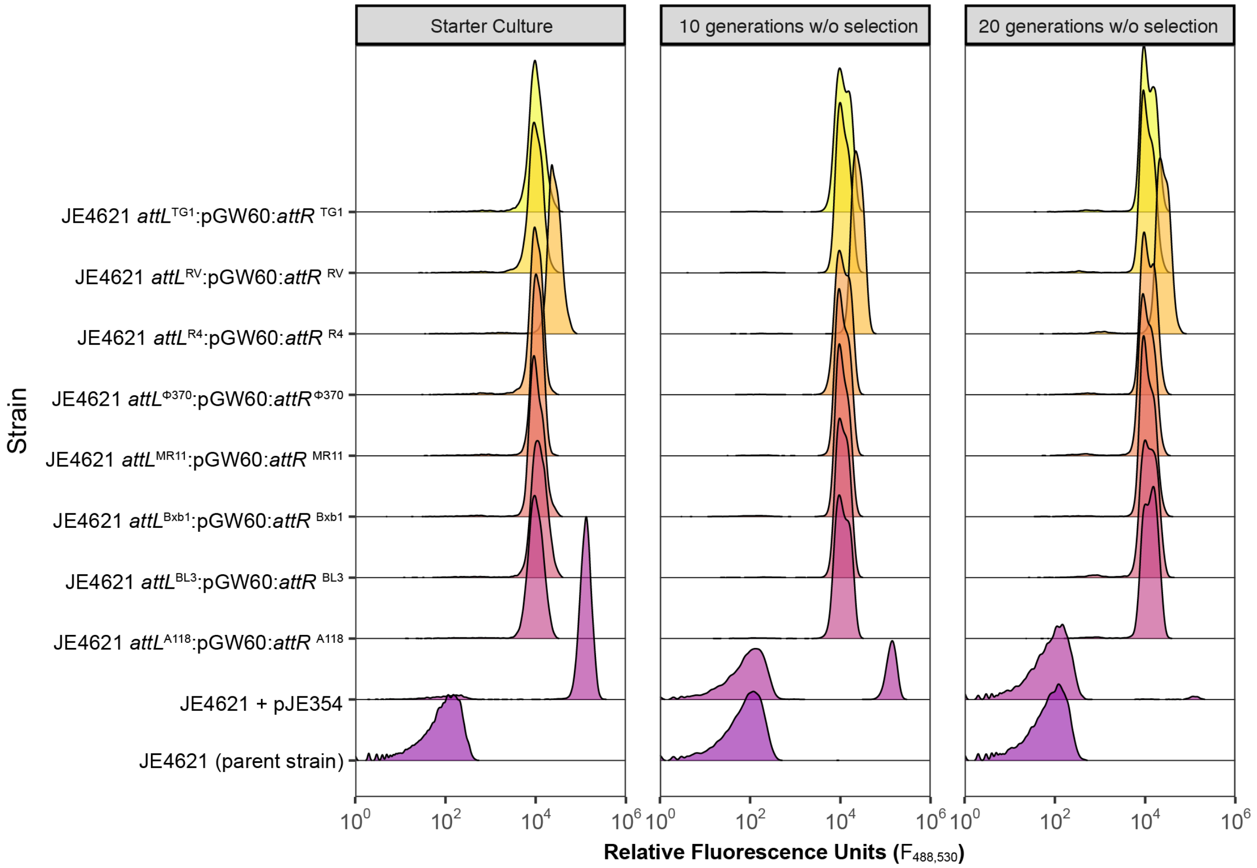


**Supplementary Fig. S3. Assays to assess stability of engineered functions in *P. fluorescens* JE4621 via SAGE or replicating plasmid.** Flow cytometry measurements of either the starter culture (green) – grown in presence of 50 µg/mL kanamycin – or cultures passaged in medium lacking antibiotic. Each passage was inoculated with a 1024-fold dilution of its precursor culture (Starter -> Passage 1 -> Passage 2) to enable 10 generations of growth prior to reaching stationary phase. The x-axis indicates the relative fluorescence ($Ex_{488}$ ,$Em_{530}$) of each cell. The Y-axis represents the fraction of cells in the culture with a given relative fluorescence value. Each plot contains data from JE4621 with no exogenous DNA, the replicating plasmid pJE354, or with pGW60 integrated using SAGE. For SAGE samples, the location of pGW60 integration is indicated.

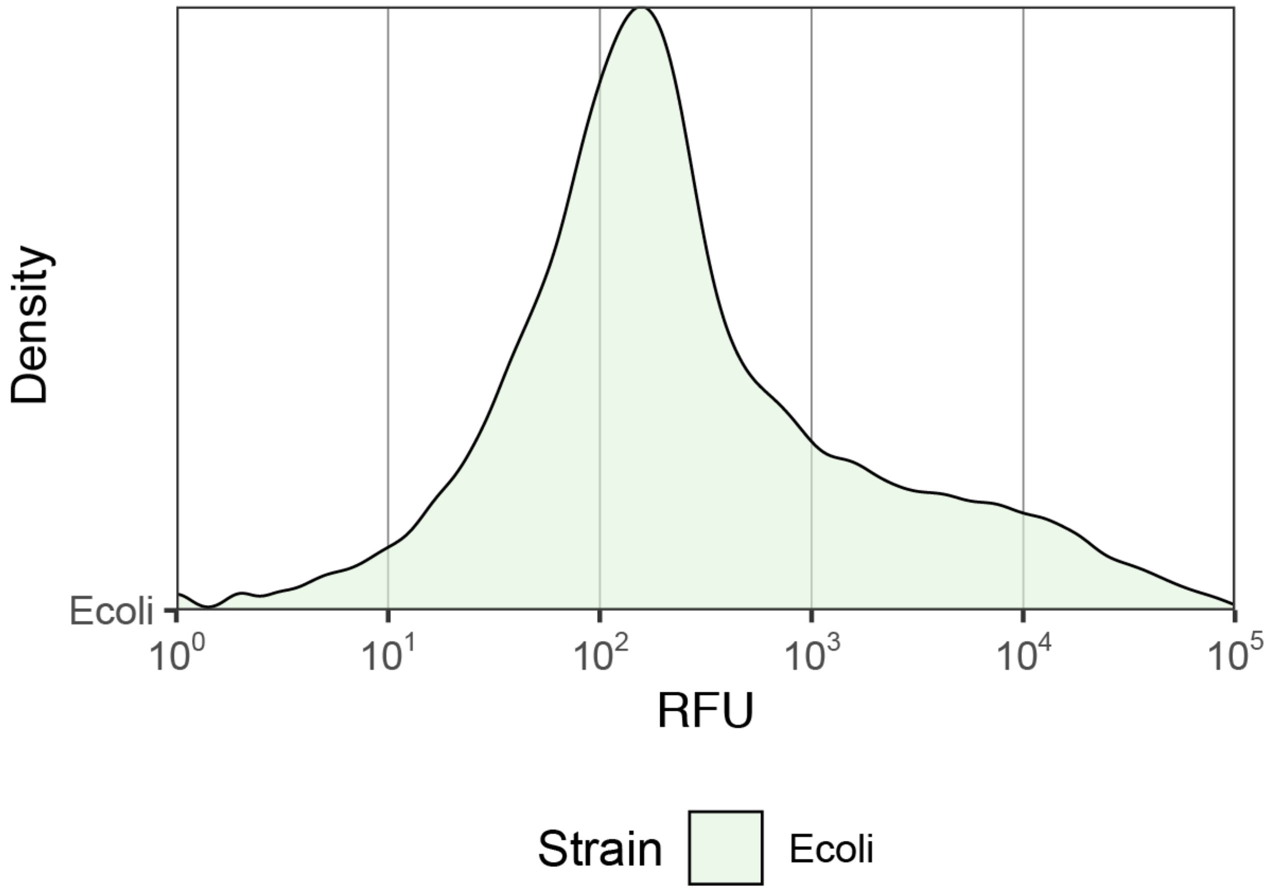


**Supplementary Fig. S4. Flow cytometry assessment of *E. coli* strain library hosting pooled promoter library plasmids.** Flow cytometry measurements of the *E. coli* plasmid library culture prior to plasmid DNA extraction. The x-axis indicates the relative fluorescence ($Ex_{488}$ ,$Em_{530}$) of each cell. Values of ~10^2 or lower are likely strains with low to no mNeonGreen expression. The Y-axis represents the fraction of cells in the culture with a given relative fluorescence value.

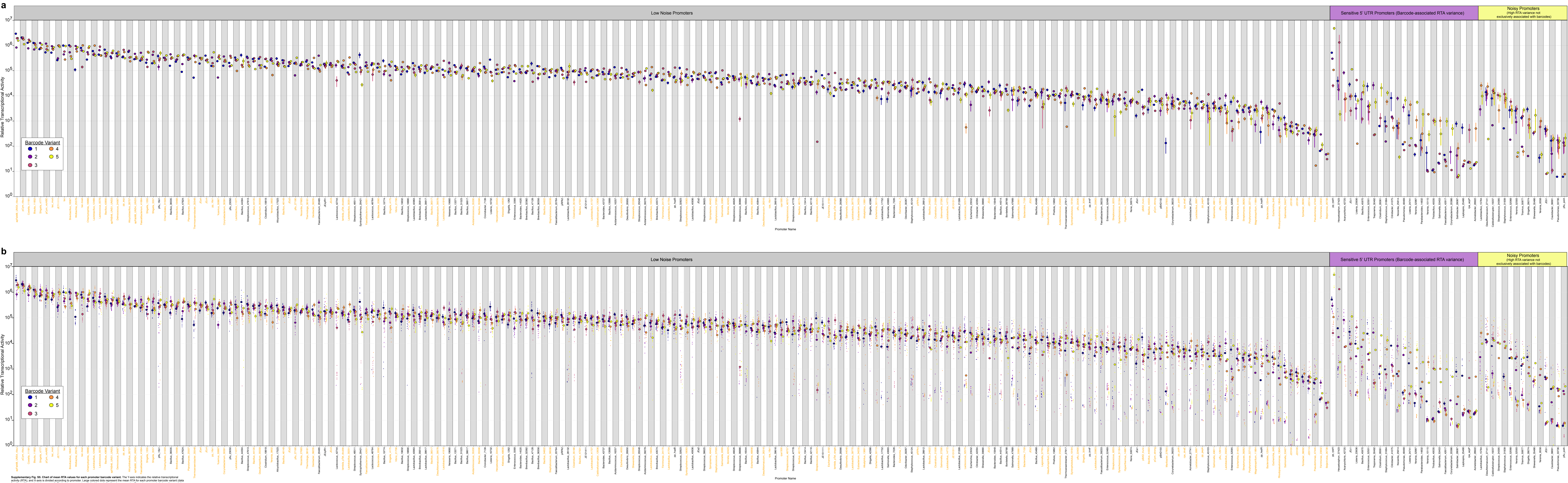


**Supplementary Fig. S5. Chart of mean RTA values for each promoter barcode variant.** The Y-axis indicates the relative transcriptional activity (RTA), and X-axis is divided according to promoter. Large colored dots represent the mean RTA for each promoter barcode variant (data for all condition variants is averaged). Each color represents a distinct barcode for the given promoter. Promoters are sorted along the x-axis by both average promoter RTA and noise classification (Low noise, Barcode-associated noise, or high noise promoters). Orange shading of promoter labels on the X-axis indicates promoters that are classified as low-noise, with condition-independent expression. (a) Error bars represent the standard error of the mean in at least 12 individual RTA measurements. (b) Small colored dots represent the each individual RTA measurement. Each color represents the associated barcode variant.

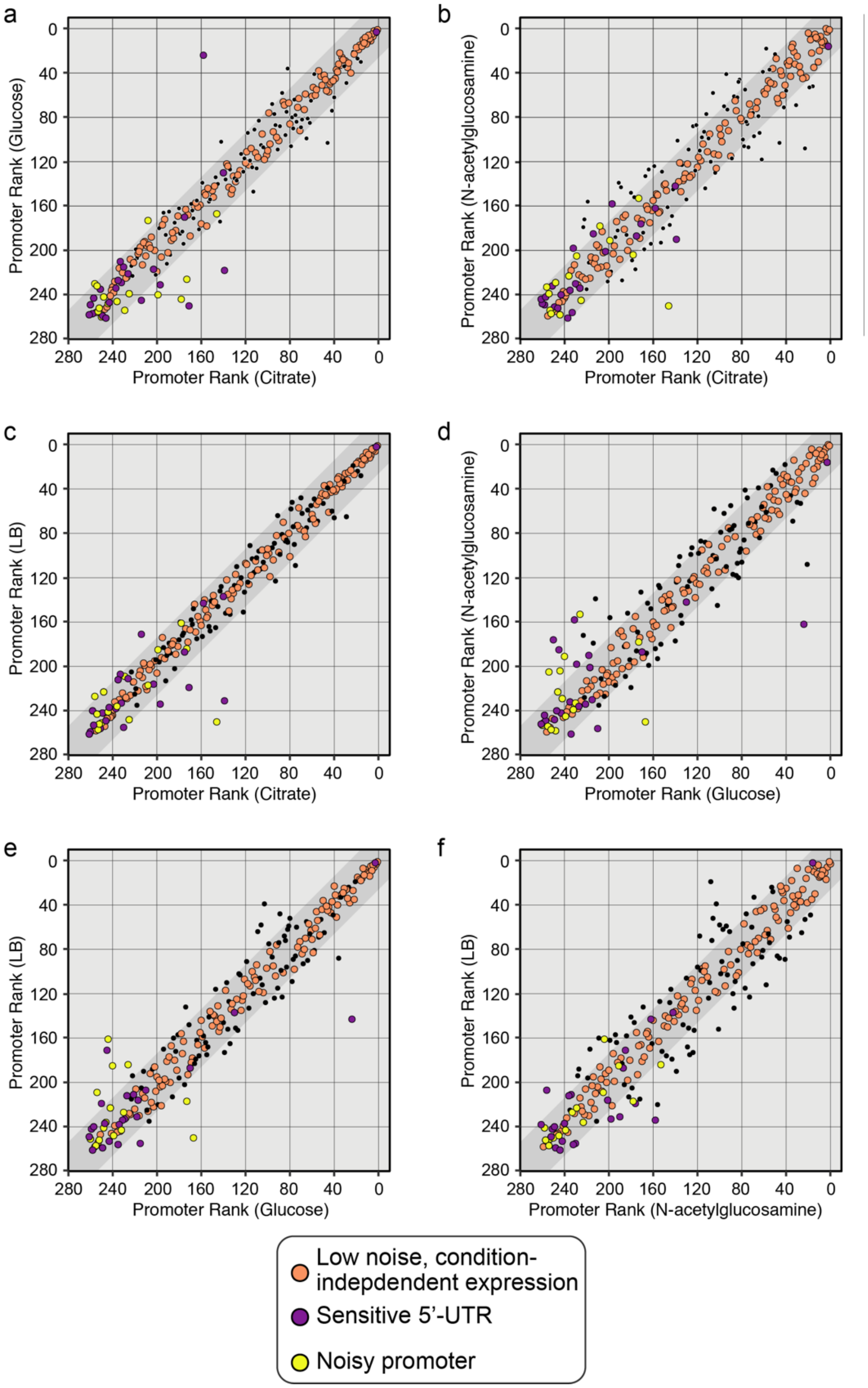


**Supplementary Fig. S6. Pairwise comparison of the relative transcriptional activity ranks for 6 combinations of conditions using all 262 promoters assayed with the *P. fluorescens* JE4621 library.** X and Y-axes represent the rank of the promoter relative to other promoters in the library when grown in the indicated condition. The dark gray band in each plot represents the range in which promoters need to fall within for all 6 pairwise conditions to be considered 'condition-independent'. All promoter rankings are determined by comparing mean of the RTAs for all barcodes and replicates of a given promoter for each condition. Orange dots represent promoters that fall within the dark gray band under all conditions, and that are also considered low noise by all other metrics. Purple dots represent promoters that have high variation in RTA values between barcode variants, and are deemed to have sensitive 5'-UTR sequences. Yellow dots represent promoters with high noise in RTA values that isn't specifically associated with barcode variation.

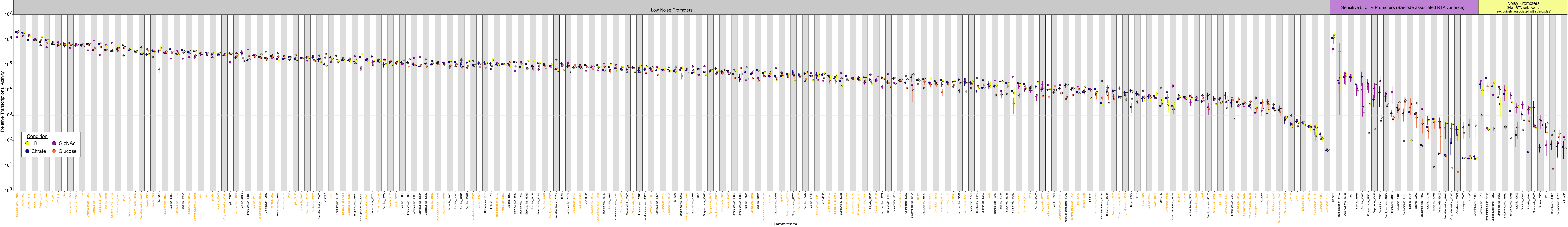


**Supplementary Fig. S7. Chart of mean RTA values for each condition.** The Y-axis indicates the relative transcriptional activity (RTA), and X-axis is divided according to promoter. Large colored dots represent the mean RTA for each condition (data for barcode variants is averaged). Each color represents a distinct condition. Promoters are sorted along the x-axis by both average promoter RTA and noise classification (Low noise, Barcode-associated noise, or high noise promoters). Orange shading of promoter labels on the X-axis indicates promoters that are classified as low-noise, with condition-independent expression. (a) Error bars represent the standard error of the mean in at least 12 individual RTA measurements.

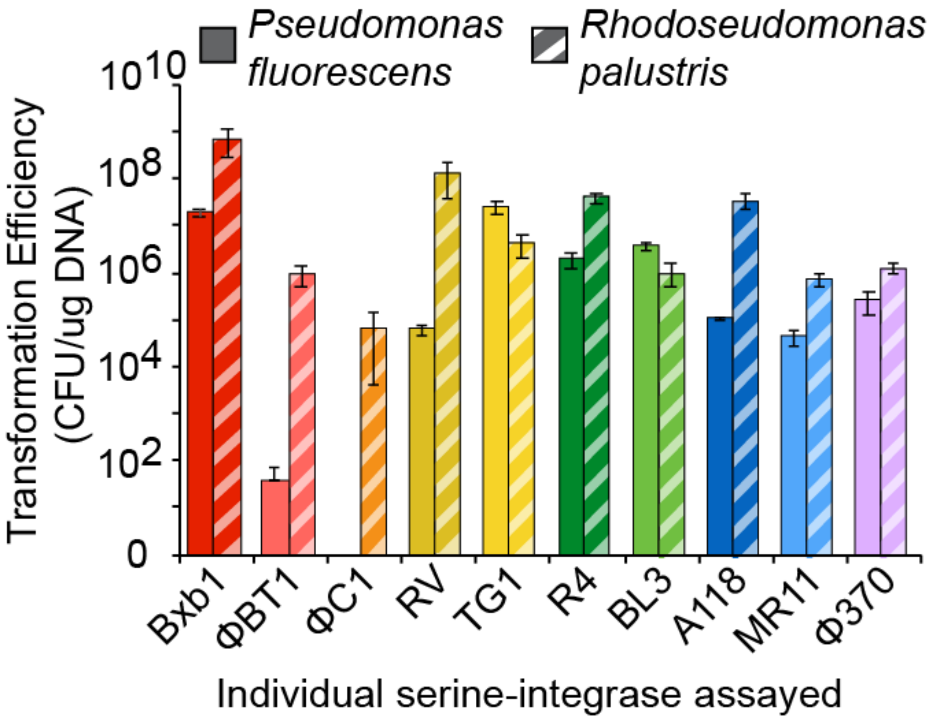


**Supplementary Fig. S8. Comparison of transformation efficiencies when using SAGE in *Pseudomonas fluorescens* SBW25 and *Rhodopseudomonas palustris* CGA009.** Y-axis represents CFU per µg of the *attP* target plasmid pGW60. All transformations were performed as co-transformations of 100 ng pGW60 with a standardized amount (250 ng) of the indicated suicide *helper* plasmid. The error bars represent standard deviation in 3 replicate experiments.

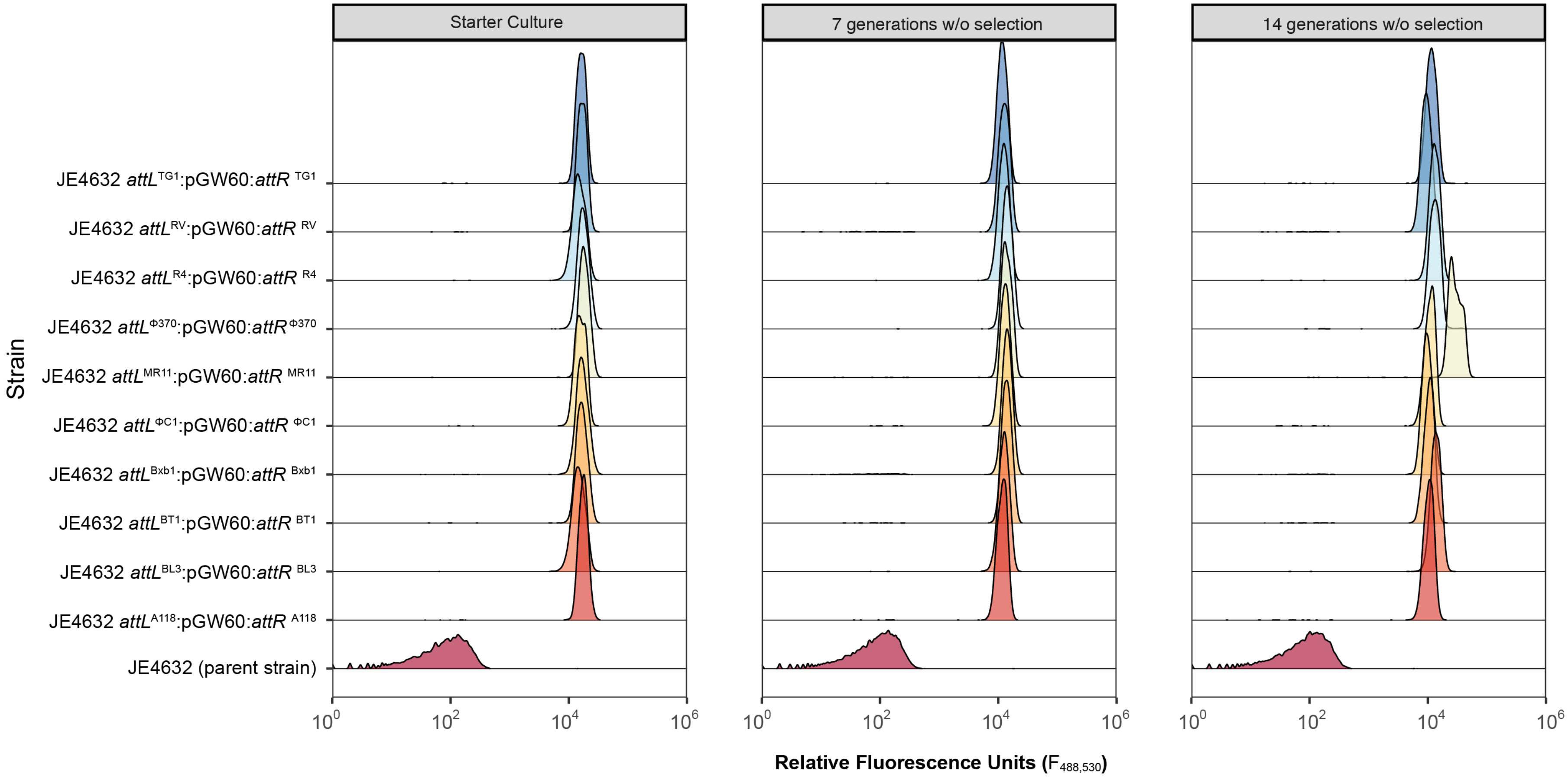


**Supplementary Fig. S9. Assays to assess stability of engineered functions in *R. palustris* JE4632 via SAGE.** Flow cytometry measurements of either the starter culture (green) – grown in presence of 200 µg/mL kanamycin – or cultures passaged in medium lacking antibiotic. Each passage was inoculated with a 128-fold dilution of its precursor culture (Starter -> Passage 1 -> Passage 2) to enable 7 generations of growth prior to reaching stationary phase. The x-axis indicates the relative fluorescence ($Ex_{488}$ ,$Em_{530}$) of each cell. The Y-axis represents the fraction of cells in the culture with a given relative fluorescence value. Each plot contains data from JE4632 with no exogenous DNA, the replicating plasmid pJE354, or with pGW60 integrated using SAGE. For SAGE samples, the location of integration is indicated.

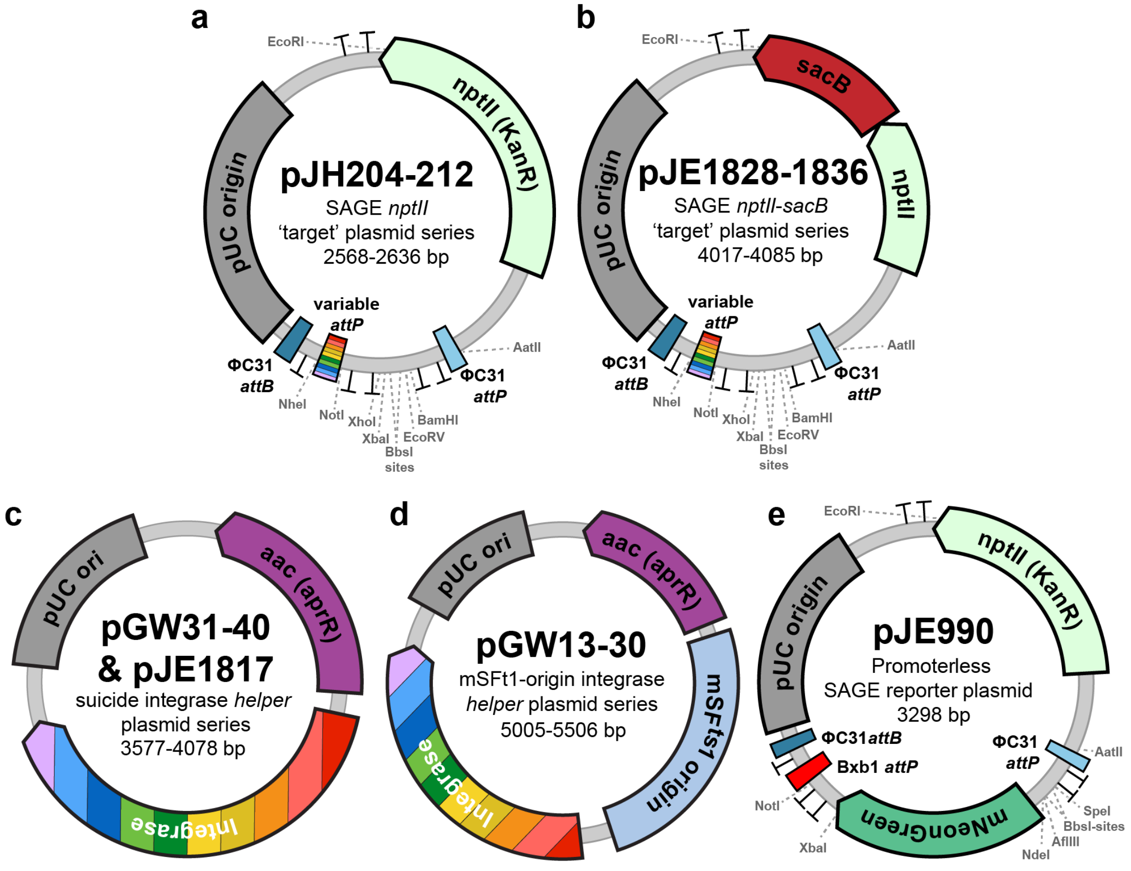


f

| Integrase | *nptII attP* 'target' plasmid | *nptII-sacB attP* 'target' plasmid | suicide integrase *helper* plasmid | mSF$^{ts1}$ ori integrase *helper* plasmid |
|---|---|---|---|---|
| Bxb1 | pJH204 | pJE1828 | pGW31 | pGW13 |
| RV | pJH205 | pJE1829 | pGW32 | pGW21 |
| Φ370 | pJH206 | pJE1830 | pGW37 | pGW20 |
| R4 | pJH207 | pJE1831 | pGW39 | pGW23 |
| ΦBT1 | pJH208 | pJE1832 | pGW33 | pGW14 |
| MR11 | pJH209 | pJE1833 | pGW36 | pGW18 |
| TG1 | pJH210 | pJE1834 | pGW38 | pGW22 |
| BL3 | pJH211 | pJE1835 | pGW40 | pGW24 |
| A118 | pJH212 | pJE1836 | pGW35 | pGW17 |
| ΦC1 | --- | --- | pGW34 | pGW16 |
| ΦC31 | --- | --- | pJE1817 | pGW30 |

**Supplementary Fig. S10**. **SAGE toolkit - plasmid collection.** Graphic plasmid maps of the SAGE plasmid collection, with the exception of poly-*attP* plasmid pGW60 (**Fig. 3**), and pJE990/GW60 *sacB*- & *pheS**-derivatives. Terminators are indicated by 'T' symbols. (**a-c**) SAGE 'target' plasmids contain an *attP* sequence to target genome integration with a specific serine integrase. Each plasmid contains a pair of ΦC31 *att* sites for plasmid backbone excision. Terminators flank both the multiple cloning site variable *attP* sequence to insulate both the integrated DNA cargo and neighboring genomic DNA (following integration). Plasmids are compatible with golden gate cloning (dual BbsI-sites) and contain restriction sites to enable modular replacement of the major components of the plasmids (selection markers, *attP*, insulating terminators, & origin of replication). Plasmid series contain either *nptII* alone, or contain a (**b**) *sacB* or (**c**) *Bacillus subtilis pheS*$^{T255S, A309G}$ counter-selection marker to enable selection for plasmid backbone excision when using a suicide ΦC31-integrase *helper* plasmid. (**d-e**) Integrase expression plasmids containing the *E. coli* origin of replication (**d**) alone or (**e**) with a temperature-sensitive mSF$^{ts1}$ origin for replication in *Pseudomonads*. (**f**) A mNeonGreen fluorescent protein reporter plasmid for testing genetic control elements (promoters, ribosomal binding sites, and terminators). Contains restriction sites for easy promoter insertion, RBS replacement, and downstream terminator replacement. (**g**) Table of SAGE plasmids described in the figure.
