## Supplementary Materials and Methods for "The SAGE genetic toolkit enables highly efficient, iterative site-specific genome engineering in bacteria"

*General culture conditions & media*

The strains and plasmids used in this study are listed in **Supplementary Table S1**. Routine cultivation of *Escherichia coli* for plasmid construction and maintenance was performed at 37 °C using LB (Lennox) medium supplemented with antibiotics (50 µg/mL kanamycin sulfate, 50 µg/mL apramycin sulfate, or 30 µg/mL gentamicin sulfate) and 15 g/L agar (for solid medium). All *Pseudomonas fluorescens* and *Rhodopseudomonas palustris* cultures were incubated at 30 °C unless otherwise indicated. Cultures were aerated with shaking at 225 rpm for test tube and shake flask cultures, and "fast shaking" 1 mM orbitals for plate reader assays.

LB (Lennox) was used for routine *Pseudomonas fluorescens* strain maintenance, competent cell preparations, starter cultures, and for plasmid stability assays. Kanamycin sulfate (50 µg/mL) or apramycin sulfate (50 µg/uL) were utilized for antibiotic selection. Van Niel's medium (ATCC medium 112) supplemented with 20 mM NaAcetate (VN-A medium) was utilized for routine cultivation of *R. palustris*. All cultivation of *R. palustris* was performed aerobically in the dark. Kanamycin sulfate (200 µg/mL) or gentamicin sulfate (µg/mL) were utilized for antibiotic selection.

Modified M9 medium (M9*) with variable nitrogen or carbon sources was utilized for shake flask experiments and fluorescent plate reader assays (47.8 mM $Na_2HPO_4$, 22 mM $KH_2PO_4$, 8.6 mM NaCl, 1 mM $MgCl_2$, 0.1 mM CaCl2, 18 µM $FeSO_4$, 1x MME trace minerals, pH adjusted to 7 with KOH). 1000x MME trace mineral stock solution contains per liter, 1 mL concentrated HCl, 0.5 g $Na_4EDTA$, 2 g $FeCl_3$, 0.05 g each $H_3BO_3$, $ZnCl_2$, $CuCl_2 \cdot 2H_2O$, $MnCl_2 \cdot 4H_2O$, $(NH_4)_2MoO_4$, $CoCl_2 \cdot 6H_2O$, $NiCl_2 \cdot 6H_2O$. For promoter testing assays M9* medium was supplemented with the following: 10 mM $NH_4Cl$ with either 10 mM glucose or 10 mM citrate, 10 mM N-acetylglucosamine, or 10 mM glucose with 10 mM urea, 10 mM sodium nitrate, or 10 mM sodium nitrite.

*Plasmid & Bacterial strain construction*

Q5® High-Fidelity DNA Polymerase (New England Biolabs - NEB) and primers synthesized by Eurofins Genomics were used in all PCR amplifications for plasmid construction. OneTaq® DNA polymerase (NEB) was used for colony PCR. EvaGreen® dye (Biotium) was supplemented at 1x final concentration in PCR reactions for qPCR applications. Plasmids were constructed by Gibson Assembly using NEBuilder® HiFi DNA Assembly Master Mix (NEB) or ligation using T4 DNA ligase (NEB). Plasmids were transformed into either competent NEB Turbo, NEB 5-alpha F'$I^q$ (NEB), Epi400 (Lucigen), or QP15 (Epi400 mated with NEB 5-alpha F'$I^q$ to transfer the mini F' plasmid to Epi400). Standard chemically competent *Escherichia coli* transformation protocols were used to construct plasmid host strains. Transformants were selected on LB (Lennox) agar plates containing antibiotics (50 µg/mL kanamycin sulfate, 50 µg/mL apramycin sulfate, or 30 µg/mL gentamicin sulfate) for selection and incubated at 37 °C. Template DNA for PCRs was either synthesized by IDT, synthesized by GenScript, or isolated from *E. coli, P. fluorescens* SBW25, or *R. palustris* CGA009 using DNeasy Blood & Tissue gDNA purification kit (Qiagen).

Zymoclean Gel DNA recovery kit (Zymo Research) was used for all DNA gel purifications. Plasmid DNA was purified from *E. coli* using GeneJet plasmid miniprep kit (ThermoScientific) or ZymoPURE II plasmid midiprep kit (Zymo Research). Sequences of plasmids were confirmed using Sanger sequencing performed by Eurofins Genomics. Plasmids used in this work are listed in **Supplementary Table S1**, and maps of plasmids constructed in this study can be found in **Supplementary File P1.zip**.

*P. fluorescens* SBW25 was used as a parent strain for all *P. fluorescens* strains in this study (**Supplementary Table S1**). Competent cells were prepared and the allelic exchange protocol used for integration of the poly-*attB* cassette was performed as described previously [1] for *P. putida* KT2440, with the exception that pK18sB [2] was utilized as the cloning vector, rather than pK18mobsacB. The same competent cell preparation and electroporation protocols above were used for electroporation of serine integrase *attP* 'target' plasmids and for replicating or non-replicating *helper* integrase expression plasmids.

*Rhodopseudomonas palustris* CGA009 was used as a parent strain for *R. palustris* strains in this study (**Supplementary Table S1**). Electrocompetent cells were prepared by pre-culturing *R. palustris* aerobically at 30 °C until the cells reached stationary phase. Fresh media was inoculated with a 1:100 dilution of the pre-culture and incubated aerobically until the cells reach mid-to-late log phase (~OD600 = 0.5 - 0.9). Once desired culture density is reached the cells were incubated on ice until chilled. All following steps were performed on ice or at 4 °C. Cells were centrifuged at 4 °C and 4000 x g for 10 minutes. Pelleted cells were washed in 10/10$^{th}$ culture volume of ice-cold 10% glycerol. The centrifugation and wash steps were performed two more times for a total of three washes. Following the final centrifugation, the cells were resuspended in 1:400$^{th}$ culture volume of ice-cold 10% glycerol and either used immediately or stored at -80 °C. Electroporation was performed as follows. 50 µL of electrocompetent cells or electrocompetent cells diluted 1:10 in ice-cold 10% glycerol were transferred to an ice-cold 0.1 cm gap electroporation cuvette. For allelic exchange, 500-1000 ng of plasmid DNA was added to cells, and the mixture was electroporated with settings at 1.75 kV, 25 µF, and 200 ohms. The same protocol was used for electroporation of serine integrase *attP* 'target' plasmids and non-replicating *helper* integrase expression plasmids. Following electroporation, 950 µL of VN-A was added to the competent cells and the resulting mixture was incubated aerobically, at 30 °C with shaking for 2 hours. Following this recovery step, various dilutions of recovery cultures were plated on selective VN-A solid medium and incubated aerobically at 30 °C. Colonies typically appear after ~5 days when *R. palustris* is grown on solid medium.

For allelic exchange in *R. palustris* we used the pJQ200SK [3] gentamicin resistance/sucrose-sensitivity selection/counter-selection method for allelic exchange with minor modifications for *R. palustris* to generate the poly-*attB* strain JE4632. For this, 700-bp regions of homology flanking RPA1300 were cloned immediately upstream and downstream of the poly-*attB* cassette within the multiple cloning site of pJQ200SK. The resulting plasmid was transformed into *R. palustris* CGA009 and resulting colonies typically arise from plasmid recombination into the genome via homologous recombination were selected by cultivation on VN-A + 200 µg/mL gentamicin sulfate. Isolated colonies were streaked for single colony isolation on VN-A medium with gentamicin. For counter-selection, we cultivated colonies from the second selective plate on VN-A + 10% sucrose. Resulting colonies have typically resulted from excision of *sacB* from the genome, as its expression caused a significant growth defect in the presence of sucrose. Resulting colonies were patched onto VN-A + 10% sucrose, and subsequently tested by colony PCR for the insertion of the poly-*attB* cassette. Positive colonies were subsequently streaked for single colony isolation again on VN-A, and resulting colonies cultivated in liquid VN-A. These final cultures were screened again by colony PCR, and correct colonies were stored as 10% glycerol stocks at -80 °C.

Primers used to screen for integration of the poly-*attP* cassette insertions in *P. fluorescens* and *R. palustris* can be found in **Supplementary Table S2**.

*Temperature-sensitive plasmid stability and pGW60 backbone excision assays*

Multiple methods were utilized to assess conditional-replication of pGW26 in *P. fluorescens* SBW25 at various temperatures. First, *P. fluorescens* was electroporated with 250 ng of pGW26 and recovered in 1 mL SOC at 25 °C for 2 hours. Dilutions of the recovery cultures were inoculated onto LB + 50 µg/mL apramycin sulfate and incubated at 25 °C or 34 °C for up to 96 hours (to confirm absence of colony formation at restrictive temperature). Colonies were enumerated and transformation efficiency determined (**Fig 2a**).

Second, we cultivated a SBW25-derivative strain containing pGW26 in LB + 50 µg/mL apramycin sulfate at 25 °C to select for plasmid maintenance. After reaching stationary phase a 1:1000 dilution of the culture was inoculated into 6 different subcultures. Subcultures were grown at either 25, 30, or 34 °C either in the presence or absence of 50 µg/mL apramycin sulfate until they reached stationary phase (typically with 24 hours). Each culture in which growth occurred was diluted to reach an $OD_{600}$ of 0.5, and further 10-fold serial dilutions were prepared. 5 µL of each dilution was spotted onto solid LB medium containing or lacking 50 µg/mL apramycin sulfate. These solid medium cultures were then incubated at 25, 30, or 34 °C for up to 72 hours, and colonies were enumerated to establish viable cell counts under each condition. Overall cell viability and % of cells maintaining replicating pGW26 within each subculture at each cultivation temperature was determined by counting colonies from 4 replicates on plates lacking and containing 50 µg/mL apramycin sulfate, respectively (**Fig 2b-c**).

Third, as a demonstration for a succinct plasmid curing protocol we performing the following plasmid curing experiment using pGW26 and pGW30 (**Fig. 4**). JE4689, which contains pGW60 integrated into the Bxb1-*attB* site of JE4621, was electroporated with either of the two aforementioned *ts*-plasmids and recovery culture was incubated at 25 °C on LB + 50 µg/mL apramycin sulfate. Twenty apramycin resistant transformants for each plasmid were streaked onto LB for single colony isolation and incubated overnight at 34 °C. A single colony from each streak was tested by colony PCR for presence of both their respective *ts*-plasmids and excision of the pGW60 plasmid backbone. The same cellular material was patched onto both LB or LB + 50 µg/mL apramycin sulfate and incubated overnight at 30 °C to confirm loss of antibiotic resistance. Primers utilized for screening plasmid loss (oPNL556/879) and excision of the pGW60 backbone from the genome of JE4689 (oPNL621/622) are listed in **Supplementary Table S2**.

*Serine integrase transformation assays*

Integrase testing assays were performed using variations of the transformation procedures described in section *Plasmid & Bacterial strain construction* with the following modifications. For assays using a *ts* plasmid to deliver the serine integrase we first transformed the *ts* plasmid into the JE4621 strain and generated competent cells hosting the *helper* plasmid. The following amounts of plasmids were electroporated into either *R. palustris* JE4632, *P. fluorescens* JE4621, *P. fluorescens* JE4624, or *P. fluorescens* JE4621 containing pGW13. Transformations included 100 ng of replicating plasmid (pJE354 or pEYF2K), 750 ng of homologous recombination plasmid (pJE1609), or 100 ng of the pGW60 *attP* 'target' plasmid that is electroporated alone or co-electroporated with 250 ng of *suicide* integrase expression *helper* plasmid. Following electroporation, cells were either resuspended in 950 µL SOC (*P. fluorescens*) or 950 µL VN-A, transferred to a microfuge tube, and incubated at 30 °C for 1 (*P. fluorescens*) or 2 (*R. palustris*) hours to allow for recovery. Due to low efficiency of homologous recombination all of the recovery volume was plated for pJE1609. For the remaining plasmids several fractions of the recovery volume were plated. Colonies were enumerated and evaluated visually for green fluorescence. For *P. fluorescens* assays, all samples were plated on LB agar supplemented with 50 µg/mL kanamycin sulfate. For *R. palustris* assays, all samples were plated on VN-A agar supplemented with 200 µg/mL kanamycin sulfate. Plates were incubated at 30 °C until colonies became visible. This was typically 24 hours for *P. fluorescens* and approximately 5 days for *R. palustris*.

Integration accuracy was determined by examining 24 colonies for each integrase by PCR to pGW60 plasmid integration into the correct locus. Colonies were cultivated with shaking (800 rpm, 1 mm orbital) at 30 °C in 96-well deep well plates using either 1 mL LB + 50 µg/mL kanamycin sulfate (*P. fluorescens*) or 1 mL VN-A + 200 µg/mL kanamycin sulfate (*R. palustris*). For *P. fluorescens* 50 µL of each culture was transferred to individual wells in a 384-well Echo source plate (Product #PP-0200 - Labcyte) for an Echo 550 Acoustic Liquid Handler (Labcyte). 500 nL of each culture was transferred into 10 µL colony PCR reactions, performed using OneTaq polymerase (New England Biolab) in a performed using a CFX-384 Touch Real-Time PCR machine (Bio-Rad). Colonies were screened using either a positive control primer set that should amplify in any *P. fluorescens* SBW25 strain (oPNL615/616) or a primer set that should only amplify a product if pGW60 has been integrated into the chromosomal poly-*attB* cassette that is downstream of *ampC* in JE4621 (oPNL629/622). Of note, oPNL629/622 generate products of distinct sizes depending on which serine integrase performs the plasmid integration. A similar approach was used in *R. palustris*, with the exception that cultures were first lysed using the following method. 200 µL of saturated culture was centrifuged at 3000 x g for 5 minutes to pellet cells. Supernatant was decanted, and cell pellets were resuspended in TE (10 mM Tris-HCl, 1 mM EDTA, pH 8) + 0.1% Triton X-100 detergent. Resuspended cells were boiled at 100 °C for 5 minutes, centrifuged again, and 50 µL of supernatant was transferred to a 384-well Echo source plate. Colonies were screened by PCR as described above using primer set oPNL819/817 to assess integration in the poly-*attB* cassette located at the RPA_1300 locus. PCR reactions that failed or otherwise demonstrated negative results for correct integration were repeated at least 1 additional time to confirm the negative result.

*Flow cytometry assessment of expression construct stability*

Starter cultures for *P. fluorescens* and *R. palustris* experiments were prepared as follows. For all wild-type strains lacking an mNeonGreen expression cassette, the strain was cultivated to stationary phase in 1 mL base media (LB for *P. fluorescens* and VN-A for *R. palustris*) lacking antibiotic. Transformant strains were generated as described in the *Serine integrase transformation assays* section above. Starter cultures for transformant strains were generated by incubating a representative colony for each strain in 1 mL kanamycin sulfate supplemented base media (50 µg/mL for *P. fluorescens* strains and 200 µg/mL for *R. palustris* strains) at 25 °C until the culture reached saturation (20 hr for *P. fluorescens,* ~60 hr for *R. palustris*). Passages were performed by diluting cultures by either 1024-fold (10 generation dilution) or 128-fold (7 generation dilution) in fresh medium and incubating at 25 °C until stationary phase was reached (~16 hours for *P. fluorescens* and ~36 hours for *R. palustris*). All cultivations were performed in 96-well deep well plates with shaking (800 rpm, 1 mm orbital). Stationary phase cultures were assessed for construct stability by flow cytometry. Briefly, stationary phase cultures were diluted 500-fold in 1x phosphate buffered saline (PBS) and fluorescence was measured using a NovoCyte Flow Cytometer (Acea Biosciences). Green fluorescence (mNeonGreen) was measured using a 488 nm laser with a 530/30 nm filtered detector. Red fluorescence (mKate2) was measured using a 561 nm laser with a 586/20 nm filtered detector. The percentage of cells considered to have lost the expression cassettes were calculated by enumeration of the fraction of cells whose fluorescence overlapped with that of the parent strain – which lacks a fluorescent protein expression cassette.

*Promoter library design, synthesis, cloning, and transformation into P. fluorescens JE4621*

A combination of 287 synthetic and natural promoter sequences were synthesized with the following parameters. Promoter sequences, barcode sequences, and promoter origin can be found in **Supplementary File T1**. Synthetic promoters were derived from commonly used promoters in the field of synthetic biology, as well as custom promoters generated for this study. Any synthetic promoters longer than 150 nt were truncated that size from the 5' end. Natural promoters were either sourced from [4], or mined from other organisms listed in **Supplementary File T1** by identifying intergenic regions upstream of genes of interest. Promoter elements for these genes were obtained by removing the sequence likely to include the ribosomal binding sequence, or specifically the 15 nt immediately upstream of the start codon of the given gene. This excision was performed to better assess promoter performance outside of the context of the native RBS, which is unlikely to be utilized if the promoter is used in synthetic expression cassettes. Sequences longer than 150 nt were trimmed to that length by removing sequence from the 5' end (distal from the start codon). For simplicity these sequences are referred to as promoters. As indicated in Figure 5a, we then generated five barcode variants for each promoter, in which a distinct 12 nt barcode sequence (Levenshtein distance of >2) was appended immediately downstream of each promoter sequence. We then added a common 18 nt spacer sequences on either end of the promoter, as well as flanking sequence containing two oppositely oriented *Bbs*I recognition sites. For promoters shorter than 150 nt, a randomly generated DNA sequence was incorporated upstream of the 5' *Bbs*I recognition site to bring the total construct sequence to 200 bp. In total, a 200-bp oligo pool of 1435 promoters was generated.

All enzymes were obtained from New England Biolabs unless otherwise noted. The promoter library was synthesized as a 1-pmol oligo mix by Twist Biosciences. The oligo library was resuspended in TE to a concentration of 5 ng/uL. 10 ng of resuspended oligo was PCR amplified for 8 cycles of PCR in a 50 µL reaction for generation of a template stock (*Pr_amp1*). All subsequent amplifications use this template as input DNA to avoid freeze-thaw cycles of the original oligo library stock. We performed a second amplification step using 3 µL of *Pr_amp1* template stock to obtain enough DNA of the library (*Pr_amp2*) for cloning into the recipient plasmid pJE990. For this we performed qPCR in 10 parallel 20 µL reactions that were stopped after the reaction exited exponential amplification stage (~7-9 cycles). All reactions used Q5 polymerase, supplemented with 1x EvaGreen (Biotium) for qPCR, and were performed using a CFX-384 Touch Real-Time PCR machine (Bio-Rad). Amplified library DNA was purified, digested with *BbsI*-HFv2, and directionally ligated into *BbsI*-linearized pJE990 with T4 DNA polymerase. Ligations were transformed into *E. coli* QP15 chemically competent cells and recovered in 1 mL SOC at 37 C for 1 hour. QP15 is an Epi400 (Lucigen) derivative containing the F'$I^Q$ plasmid from NEB 5-$\alpha$ F'$I^Q$ (transferred by conjugation). A 10-µL aliquot of the recovery mixture was diluted and plated on LB + 50 µg/mL kanamycin sulfate to determine the cloning efficiency and library coverage, and the remaining 990 µL was propagated through two subsequent overnight liquid selections using 25 mL and then 150 mL of LB-Lennox (BD Biosciences) + 50 µg/mL kanamycin sulfate grown at 30 °C, 200 rpm. The latter culture was inoculated with 1 mL of the 25 mL culture. The pJE990-derived library, referred to as pLibrary (**Fig 5a**) was cloned with >50× coverage as determined by dividing the number of colony-forming units by the size of the designed library. Plasmid DNA was then extracted from library cultures with a ZymoPURE II Midiprep kit (Zymo Research) for subsequent transformation into final the host strains.

The plasmid pool pLibrary was integrated into the Bxb1 *attB* site in the genome of *P. fluorescens* JE4621 and JE4624 by co-electroporation of 200 ng pLibrary with 3 µg of pGW31 into 50 µL of electrocompetent competent cells that had been diluted 1:10 in 10% glycerol. The electroporated cells were resuspended in 950 µL SOC, incubated at 30 °C for 1 hour with shaking. As above, dilutions generated from 10 µL of the recovery culture were plated onto LB supplemented with 50 µg/mL kanamycin sulfate, and resulting colonies were quantified to identify the size of the library. The remaining 990 µL of the recovery culture was inoculated into 50 mL LB + 50 µg/mL kanamycin sulfate and incubated overnight at 30 °C. This culture was diluted with LB to an $OD_{600}$ of 1. 1 mL of the diluted culture was inoculated into 200 mL LB + 50 µg/mL kanamycin sulfate and incubated overnight at 25 °C with 200 rpm to yield the final *P. fluorescens* JE4621 and JE4624 libraries. The final JE4621 library contained ~400,000 transformants and the JE4624 library contained ~140,000 transformants. Glycerol stocks of the library cultures in final host strains were made after the second passage in liquid selection. The stocks contained 1 mL of culture diluted to $OD_{600}$ = 1. These stocks were used for all subsequent experiments. The production of mNeonGreen by members of the *E. coli* and *P. fluorescens* libraries were evaluated qualitatively by flow cytometry with a Novocyte Flow Cytometer (Acea Biosciences). Final library cultures were diluted 1:1000 in PBS and 125,000 events were analyzed for fluorescence using a 488 nm laser with a 530/30 nm filtered detector.

*Library growth, DNA-seq and RNA-seq*

Starter cultures for assays with JE4621 were generated by inoculating an entire 1 mL $OD_{600}$ glycerol stock into 100 mL LB supplemented with 50 µg/mL kanamycin sulfate and incubating the culture with 250 rpm shaking at 30 °C until the culture reached an $OD_{600}$ > 1. The recovered culture was washed twice in 100 mL 1x M9* medium to remove residual LB medium. Starter cultures for the assays were prepared by inoculating 1 mL of the washed cells into 100 mL of each of the following media and incubated overnight with 250 rpm shaking at 30 C: LB, M9* + 20 mM glucose, M9* + 20 mM sodium citrate, and M9* + N-acetylglucosamine. Four replicate assay cultures for each media type were prepared. Library measurements were carried out 100 mL cultures inoculated to an $OD_{600}$ = 0.01 from starter cultures grown in the same experimental medium. These cultures were grown to an $OD_{600}$ of 0.2 to 0.4 at 30 °C with shaking at 250 rpm. Upon reaching the target density cultures were rapidly aliquoted into 2x 25 mL aliquots and a 50 mL aliquot, then harvested by centrifugation at 7000x g for 3 minutes at 4 °C. Culture supernatants were decanted and cell pellets flash frozen in liquid nitrogen prior to storage at -80 °C. Frozen pellet from the 25 mL aliquots were later resuspended in 750 µL DNA/RNA Shield (Zymo Research) and both total RNA and genomic DNA were extracted with the ZymoBIOMICs DNA/RNA miniprep kit (ZymoResearch) according to manufacturer's parallel purification protocol.

In brief, the RNA sequencing library was prepared by reverse transcription and common adaptor ligation at the 3'-end of the cDNA. The RNA-seq and DNA-seq libraries were prepared as described by *Yim et al.* [4] with minor modifications as described below. Primers and adapter sequences for each step are found in **Supplementary Table S2**. First, ProtoScript II First Strand cDNA Synthesis Kit (New England Biolabs) was used rather than Maxima reverse transcriptase (Thermo Scientific). Accordingly, the cycling conditions were modified to: 42°C for 60 minutes, followed by 10 cycles of 50°C for 2 minutes and 42°C for 2 minutes, with a final 5 minutes at 80°C to deactivate the enzyme. Illumina indexes and adaptors were added to both cDNA and input library DNA using a two-step amplification process. Second, qPCRs to attach Illumina indices and adaptors used the DNA-binding dye EvaGreen (Biotium) rather than SYBR Green I (Invitrogen), and templates for the first step PCR were as follows: 1 µL undiluted cDNA library template with 3' adapter, and 300 ng of gDNA. First step PCRs (Amp1) were performed separately for RNA-seq and DNA-seq libraries, as they required different numbers of cycles to reach exponential amplification. Amplification was stopped as soon as exponential phase ceased, typically around 20 cycles for cDNAs and around 15 cycles for gDNA. Samples were diluted 1:100 and amplified again using the same protocol using indexing primers for 10 cycles (Amp2). Samples were then co-purified and examined on a 2% agarose gel to verify correct band sizes of ~ 200 bp for cDNA and 350 bp for input plasmid DNA libraries.

*Quantifying relative transcriptional activity from high-throughput sequencing*

Using custom Python scripts adapted from Yim *et al.* [4] (https://github.com/ssyim/DRAFTS), we mapped DNA and RNA reads to promoter and 12-nt barcode sequences. To account for a range of promoter lengths in our library, we adjusted acceptable promoter length thresholds in the DNA and RNA processing scripts appropriately. For each replicate, we used a 5-read cutoff for DNA samples and did not calculate the transcription rate for barcodes with fewer than 5 DNA reads. For each replicate sample, we calculated the transcription rate $T_i$ of each promoter barcode variant $i$ with RNA counts $R_i$ and DNA counts $D_i$ as $T_i = \frac{(R_i)/\sum_j R_j}{D_i/\sum_j D_j}$. When calculating transcription rates, the normalized RNA count ($(R_i)/\sum_j R_j$) for barcodes with no RNA reads was set to half of the minimum, non-zero normalized RNA count for that replicate. Transcription rates were then normalized to a minimum of 1 by dividing all transcription rates by the minimum value. Replicate data point $a$ (within condition $l$ and barcode variant $i$) from group $X$ was considered an outlier, and discarded from further analysis if its value was $> 3(MAD)$, or mean absolute deviation, from the median $\bar{X}$ and subsequently discarded. MAD was calculated as $median(|X_a - \bar{X}|)$. Following this processing, any promoter barcode variants that lacked at least 3 replicate data points for each condition was discarded. Following these filtering steps, any promoter lacking at least 3 barcode variants was discarded from further analysis. The resulting data is found in **Supplementary File T3**.

*Analyzing promoter strength, noise, and condition-independence from high-throughput sequencing*

Overall promoter strength ($PS_k$) was calculated using $\frac{1}{n}(\sum T_k)$ ,using transcription rate $T_k$ of each promoter $k$ and $n$ replicate data points. For figure 5d, the error bars represent the standard error of the mean ($se_k$) for $PS_k$. Barcode promoter strength ($PS_i$) was calculated using $\frac{1}{n}(\sum T_i)$ , using transcription rate $T_i$ of each promoter barcode variant $i$ and $n$ replicate data points. For figure 5c and Supplementary Fig. 5, the error bars for barcode variants represent the standard error of the mean ($se_i$) for $PS_i$. Barcode-associated noise (or variance) was calculated as follows. Overall promoter strength ($PS_k$) was calculated by an alternative method $\frac{1}{n}(\sum PS_i)$ that produces the same value as $\frac{1}{n}(\sum T_k)$, but the standard error of the mean with this method ($se_{ik}$) provides information on the variance in promoter strength between barcode variants. $\%se_{ik}$ was calculated as $\%se_{ik} = se_{ik}/PS_k$. Promoters were considered to have a sensitive 5'-UTR (high barcode noise / variance) if $\%se_{ik} > 3\big(median(\%se_{ik})\big)$. Non-barcode-associated noise was calculated as follows. $\%se_k$ was calculated as $\%se_k = se_k/PS_k$. Promoters were considered to be "generally" highly noisy if $\%se_k > 3\big(median(\%se_k)\big)$ and the promoter was not also considered to have a sensitive 5'-UTR that explains the noise.

Promoter condition-independence was determined as follows. Promoter strength ($PS_l$) within a given condition was calculated using $\frac{1}{n}\big(\sum T_{k,l}\big)$, using transcription rate $T_{k,l}$ of each promoter barcode variant $k$ in condition $l$, and with $n$ replicate data points. Error bars for condition variants in Fig. 5c and Supplementary Figure 7 represent the standard error of the mean ($se_l$) for $PS_l$. For each condition (LB, M9 + glucose, M9 + citrate, M9 + N-acetylglucosamine) the rank ($R_l$) of the promoter from strongest (1) to weakest (262) was determined. A promoter was considered to be condition independent if $\big|R_{l(condition\ a)} - R_{l(condition\ b)}\big|$ for all pairwise comparisons between conditions is never $> 1/10(n)$, where $n$ is the number of promoters that passed the initial filtering step. Promoters considered "low noise, and condition-independent" are promoters which are not considered to be noisy (barcode-associated or non-barcode-associated) and are also considered to be condition-independent.

*Plate reader promoter testing assays*

Reporter strains were constructed by transformation of *P. fluorescens* JE4621 with a combination of reporter plasmid and pGW31 (Bxb1 integrase *suicide helper* plasmid), using the methods described in section *Serine integrase transformation assays*. Reporter plasmids are listed in **Supplementary Table S1**. Strains were initially cultivated overnight at 30 °C, 225 rpm in either 5 mL LB (JE4621) or 5 mL LB + 50 µg/mL kanamycin sulfate (JE4621 transformants). 5 mL starter cultures in M9* + 10 mM glucose + 10 mM NH4Cl were inoculated with 1% of the initial culture and similarly incubated. Coupled growth and fluorescence assays were performed with a Neo2SM (Bio-Tek) plate reader using 60 µL/well of M9* medium supplemented with the following carbon and nitrogen sources: 10 mM ammonium chloride and either 10 mM glucose, 10 mM citrate, or 10 mM N-acetylglucoseamine, or 10 mM glucose and either 10 mM urea, 10 mM sodium nitrate, or 10 mM sodium nitrite. Assays were performed using black-walled, µClear, flat-bottom, 384-well plates (Greiner Bio-One) with a BreatheEasy seal (USA Scientific). Plate cultures were inoculated using an Echo Acoustic Liquid Handler with 0.1% inoculum from starter cultures, and incubated overnight in the Neo2SM at 30 °C, with the fast shaking setting, 1 mm orbital. OD600 and fluorescence ($F_{501,520}$ for mNeonGreen) measured every 12 minutes. Reporter expression per cell was estimated by dividing relative fluorescence units (RFU) by $OD_{600}$ (as a proxy for cell number) for each time point within mid-log growth phase ($OD_{600}$ = 0.089-0.16) and averaging those values for each sample. Background absorbance and fluorescence readings from wells containing media blanks were averaged and subtracted from sample readings prior to analysis. Background autofluorescence not associated with mNeonGreen expression was subtracted by subtracting the mean RFU / $OD_{600}$ value from the parent strain (JE4621) from each sample. Figure 5f and 5g relative promoter strengths were derived by averaging the mean RFU/$OD_{600}$ values for each condition shown in the panel to generate a single RFU/$OD_{600}$ value for each promoter. These values were then normalize by dividing each value by the RFU/$OD_{600}$ of JEc2 (performed independently for each panel). The resulting relative promoter strength values were then multiplied by 10 to enable simpler comparison with weaker promoters. Error bars represent the standard deviation in 3 replicates and standard deviations are two-sided.

*Flow cytometry assessment of relative promoter strength*

Reporter strains were constructed by transformation of *R. palustris* JE4632 with a combination of reporter plasmid and pGW31 (Bxb1 integrase *suicide helper* plasmid), using the methods described in section *Serine integrase transformation assays*. Reporter plasmids are listed in **Supplementary Table S1**. Strains were initially cultivated overnight at 30 °C, 225 rpm in either 5 mL VN-A (JE4632) or 5 mL VN-A + 200 µg/mL kanamycin sulfate (JE4632 transformants). Stationary phase cultures were diluted 1:500 in PBS, and analyzed by flow cytometry using a NovoCyte Flow Cytometer (Acea Biosciences). Green fluorescence (mNeonGreen) was measured using a 488 nm laser with a 530/30 nm filtered detector. Median green fluorescence values for each strain were used to calculate relative promoter strength. Autofluorescence background and fluorescence associated with read-through transcription of mNeonGreen were accounted for by subtracting the median green fluorescence of the strain hosting a promoterless reporter plasmid (pJE990) from each sample prior to calculating the relative promoter strength. This allows measurement of the contribution of each promoter towards production of mNeonGreen.

| **Supplementary Table S1. Strains and plasmids used in this work** | | |
|---|---|---|
| **Name** | **Relevant Genotype** | **Source** |
| *Strains* | | |
| NEB 5-alpha F'Iq | *Escherichia coli* F´ *proA*$^{+}$*B*$^{+}$ *lacI*$^{q}$ *Δ(lacZ)M15 zzf::Tn10* (Tet$^{R}$) / *fhuA2Δ(argF-lacZ)U169 phoA glnV44 Φ80Δ(lacZ)M15 gyrA96 recA1 relA1 endA1 thi-1 hsdR17* | New England Biolabs |
| Epi400 | *Escherichia coli* F$^{-}$ *mcrA Δ(mrr-hsdRMS-mcrBC) Φ80dlacZΔM15 ΔlacX74 recA1 endA1 araD139 Δ(ara, leu)7697 galU galK λ*$^{-}$ *rpsL (Str*$^{R}$*) nupG trfA tonA pcnB4 dhfr* | Lucigen |
| QP15 | *Escherichia coli* F´ *proA*$^{+}$*B*$^{+}$ *lacI*$^{q}$ *Δ(lacZ)M15 zzf::Tn10* (Tet$^{R}$) */ mcrA Δ(mrr-hsdRMS-mcrBC) Φ80dlacZΔM15 ΔlacX74 recA1 endA1 araD139 Δ(ara, leu)7697 galU galK λ*$^{-}$ *rpsL (Str*$^{R}$*) nupG trfA tonA pcnB4 dhfr* | this work |
| SBW25 | *Pseudomonas fluorescens* SBW25 | [5] |
| CGA009 | *Rhodopseudomonas palustris* CGA009 | [6] |
| JE4621 | *P. fluorescens* SBW25 *ampC:10x poly-attB* | this work |
| JE4624 | *P. fluorescens* SBW25 PFLU5798*:10x poly-attB* | this work |
| JE4632 | *R. palustris* CGA009 ΔRPA1300:*10x poly-attB* | this work |
| JE4689 | *P. fluorescens* JE4621 *attL*$^{Bxb1}$:pGW60: *attR*$^{Bxb1}$ | this work |
| JE4670 | QP15 + pLibrary | this work |
| JE4671 | *P. fluorescens* JE4621 *attL*$^{Bxb1}$:pLibrary: *attR*$^{Bxb1}$ | this work |
| JE4672 | *P. fluorescens* JE4624 *attL*$^{Bxb1}$:pLibrary: *attR*$^{Bxb1}$ | this work |
| JE4762 | *P. fluorescens* JE4621 *attL*$^{Bxb1}$:pJE990: *attR*$^{Bxb1}$ | this work |
| JE4763 | *P. fluorescens* JE4621 *attL*$^{Bxb1}$:pJE1045: *attR*$^{Bxb1}$ | this work |
| JE4764 | *P. fluorescens* JE4621 *attL*$^{Bxb1}$:pJE1046: *attR*$^{Bxb1}$ | this work |
| JE4765 | *P. fluorescens* JE4621 *attL*$^{Bxb1}$:pJE1047: *attR*$^{Bxb1}$ | this work |

| | | |
|---|---|---|
| JE4766 | *P. fluorescens* JE4621 *attL*$^{Bxb1}$:pJE1279:*attR*$^{Bxb1}$ | this work |
| JE4767 | *P. fluorescens* JE4621 *attL*$^{Bxb1}$:pJE1280:*attR*$^{Bxb1}$ | this work |
| JE4768 | *P. fluorescens* JE4621 *attL*$^{Bxb1}$:pJE1281:*attR*$^{Bxb1}$ | this work |
| JE4769 | *P. fluorescens* JE4621 *attL*$^{Bxb1}$:pJE1282:*attR*$^{Bxb1}$ | this work |
| JE4770 | *P. fluorescens* JE4621 *attL*$^{Bxb1}$:pJE1283:*attR*$^{Bxb1}$ | this work |
| JE4771 | *P. fluorescens* JE4621 *attL*$^{Bxb1}$:pJE1284:*attR*$^{Bxb1}$ | this work |
| JE4772 | *P. fluorescens* JE4621 *attL*$^{Bxb1}$:pJE1285:*attR*$^{Bxb1}$ | this work |
| JE4773 | *P. fluorescens* JE4621 *attL*$^{Bxb1}$:pJE1286:*attR*$^{Bxb1}$ | this work |
| **Name** | **Relevant Information** | **Source** |
| *Plasmids* | | |
| pK18sB | pUC origin, *nptII*, *sacB* | [7] |
| pJQ200SK | p15a origin, *sacB*, GmR, *mob* | [3] |
| pJE354 | pBBR1 origin, *aphI*, P$_{tac}$:*mNeonGreen* | this work |
| pEYF2K | pBBR1 origin, *nptII*, *mKate2*, *mob*, *lacZalpha* | unpublished |
| pJE1609 | pJQ200SK ΔpRPA3 | this work |
| pJE1610 | pJQ200SK ΔRPA1300::*10x poly-attB* | this work |
| pJE1700 | pK18sB *ampC*(PFLU3467):*10x poly-attB* | this work |
| pJE1701 | pK18sB PFLU5798:*10x poly-attB* | this work |
| pGW26 | pUC origin, mSF$^{ts1}$, *aac* (AprR) | this work |
| pGW13 | pGW26 P$_{tac}$:*Bxb1 integrase* | this work |
| pGW14 | pGW26 P$_{tac}$:ϕ*BT1 integrase* | this work |
| pGW16 | pGW26 P$_{tac}$:ϕC*1 integrase* | this work |
| pGW17 | pGW26 P$_{tac}$:A118 *integrase* | this work |
| pGW18 | pGW26 P$_{tac}$:MR1*1 integrase* | this work |
| pGW20 | pGW26 P$_{tac}$:ϕ370 *integrase* | this work |
| pGW21 | pGW26 P$_{tac}$:RV *integrase* | this work |
| pGW22 | pGW26 P$_{tac}$:TG*1 integrase* | this work |
| pGW23 | pGW26 P$_{tac}$:R4 *integrase* | this work |
| pGW24 | pGW26 P$_{tac}$:*BL3 integrase* | this work |

| | | |
|---|---|---|
| pGW30 | pGW26 $P_{tac}$:ϕC31 *integrase* | this work |
| pGW31 | pGW26 $P_{tac}$:*Bxb1 integrase* ΔmSF$^{ts1}$ | this work |
| pGW32 | pGW26 $P_{tac}$:RV *integrase* ΔmSF$^{ts1}$ | this work |
| pGW33 | pGW26 $P_{tac}$:ϕ*BT1 integrase* ΔmSF$^{ts1}$ | this work |
| pGW34 | pGW26 $P_{tac}$:ϕC*1 integrase* ΔmSF$^{ts1}$ | this work |
| pGW35 | pGW26 $P_{tac}$:A118 *integrase* ΔmSF$^{ts1}$ | this work |
| pGW36 | pGW26 $P_{tac}$:MR1*1 integrase* ΔmSF$^{ts1}$ | this work |
| pGW37 | pGW26 $P_{tac}$:ϕ370 *integrase* ΔmSF$^{ts1}$ | this work |
| pGW38 | pGW26 $P_{tac}$:TG*1 integrase* ΔmSF$^{ts1}$ | this work |
| pGW39 | pGW26 $P_{tac}$:R4 *integrase* ΔmSF$^{ts1}$ | this work |
| pGW40 | pGW26 $P_{tac}$:*BL3 integrase* ΔmSF$^{ts1}$ | this work |
| pJE1817 | pGW26 $P_{tac}$:ϕC31 *integrase* ΔmSF$^{ts1}$ | this work |
| pJE990 | pUC origin, *nptII*, *mNeonGreen* (promoterless), Bxb1 *attP* | [8] |
| pJE1045 | pJE990 $P_{tac}$:*mNeonGreen* | this work |
| pJE1046 | pJE990 $P_{JE1312}$:*mNeonGreen* | this work |
| pJE1047 | pJE990 $P_{JE1611}$:*mNeonGreen* | this work |
| pJE1279 | pJE990 $P_{JEb1}$:*mNeonGreen* | this work |
| pJE1280 | pJE990 $P_{JEc1}$:*mNeonGreen* | this work |
| pJE1281 | pJE990 $P_{JEa1}$:*mNeonGreen* | this work |
| pJE1282 | pJE990 $P_{JEc2}$:*mNeonGreen* | this work |
| pJE1283 | pJE990 $P_{JEb2}$:*mNeonGreen* | this work |
| pJE1284 | pJE990 $P_{JEa2}$:*mNeonGreen* | this work |
| pJE1285 | pJE990 $P_{JEc3}$:*mNeonGreen* | this work |
| pJE1286 | pJE990 $P_{JEa3}$:*mNeonGreen* | this work |
| pLibrary | pJE990 $P_{library}$:*mNeonGreen* | this work |
| pGW60 | pJE990 $P_{tac}$:*mNeonGreen,* 10x poly-*attP* cassette | this work |
| pJH204 | pUC origin, *nptII*, Bxb1 *attP* | this work |
| pJH205 | pUC origin, *nptII*, RV *attP* | this work |
| pJH206 | pUC origin, *nptII*, ϕ370 *attP* | this work |
| pJH207 | pUC origin, *nptII*, R4 *attP* | this work |
| pJH208 | pUC origin, *nptII*, ϕBT1 *attP* | this work |
| pJH209 | pUC origin, *nptII*, MR11 *attP* | this work |
| pJH210 | pUC origin, *nptII*, TG1 *attP* | this work |
| pJH211 | pUC origin, *nptII*, BL3 *attP* | this work |

| | | |
|---|---|---|
| pJH212 | pUC origin, *nptII*, A118 *attP* | this work |
| pJE1828 | pUC origin, *nptII-sacB*, Bxb1 *attP* | this work |
| pJE1829 | pUC origin, *nptII-sacB*, RV *attP* | this work |
| pJE1830 | pUC origin, *nptII-sacB*, ϕ370 *attP* | this work |
| pJE1831 | pUC origin, *nptII-sacB*, R4 *attP* | this work |
| pJE1832 | pUC origin, *nptII-sacB*, ϕBT1 *attP* | this work |
| pJE1833 | pUC origin, *nptII-sacB*, MR11 *attP* | this work |
| pJE1834 | pUC origin, *nptII-sacB*, TG1 *attP* | this work |
| pJE1835 | pUC origin, *nptII-sacB*, BL3 *attP* | this work |
| pJE1836 | pUC origin, *nptII-sacB*, A118 *attP* | this work |

| Suppplementary Table S2. Oligos | | |
|---|---|---|
| **Oligo** | **Oligo Sequence (5'->3')** | **Purpose** |
| *Promoter library amplification* | | |
| oPNL824-prom_lib_fwd2 | aacgtaccgagGAAGACaaGTCTgactgctgcgagtc | forward primer for amplification of P. fluorescens promoter library from twist biosciences for cloning into pJE990 |
| oPNL825-prom_lib_rev2 | ttcttgcttGAAGACcataagCTTAcGAGACCggacca c | reverse primer for amplification of P. fluorescens promoter library from twist biosciences for cloning into pJE990 |
| *Illumina sequencing library prep* | | |
| oPNL887-mNeon-RT | TGCAGTTCATGCGTAGCTGGCAAC | Reverse transscription from mNeonGreen |
| oPNL888-3'-adaptor | /5Phos/ NNATGTACTCTGCGTTGATACCACTGCTT /3SpC3/ | Adaptor for 3'-cDNA end ligation |
| oPNL889-DNA-amp1-F | GAGTTCAGACGTGTGCTCTTCCGATCTGTCTGA CTGCTGCGAGTC | Fwd Primer for 1st amplification of DNA |
| oPNL890-RNA-amp1-F | GAGTTCAGACGTGTGCTCTTCCGATCTAAGCA GTGGTATCAACGC | Fwd Primer for 1st amplification of RNA |
| oPNL891-Amp1-R-N3 | CCTACACGACGCTCTTCCGATCT NNN CCTGTGTGAGTTAATCTTAAGCT | Rev primer for 1st amplification of DNA and RNA |
| oPNL892-Amp1-R-N4 | CCTACACGACGCTCTTCCGATCT NNNN CCTGTGTGAGTTAATCTTAAGCT | Rev primer for 1st amplification of DNA and RNA |
| oPNL893-Amp1-R-N5 | CCTACACGACGCTCTTCCGATCT NNNNN CCTGTGTGAGTTAATCTTAAGCT | Rev primer for 1st amplification of DNA and RNA |
| oPNL894-Amp1-R-N6 | CCTACACGACGCTCTTCCGATCT NNNNNN CCTGTGTGAGTTAATCTTAAGCT | Rev primer for 1st amplification of DNA and RNA |
| oPNL895-Amp2-F-IT058 | CAAGCAGAAGACGGCATACGAGAT ACAAAC GTGACTGGAGTTCAGACGTGTGCTCTTC | Fwd primer for 2nd amplification (P7) |
| oPNL896-Amp2-F-IT008 | CAAGCAGAAGACGGCATACGAGAT ACTTGA GTGACTGGAGTTCAGACGTGTGCTCTTC | Fwd primer for 2nd amplification (P7) |
| oPNL897-Amp2-F-IT030 | CAAGCAGAAGACGGCATACGAGAT CACCGG GTGACTGGAGTTCAGACGTGTGCTCTTC | Fwd primer for 2nd amplification (P7) |
| oPNL898-Amp2-F-IT020 | CAAGCAGAAGACGGCATACGAGAT GTGGCC GTGACTGGAGTTCAGACGTGTGCTCTTC | Fwd primer for 2nd amplification (P7) |
| oPNL899-Amp2-F-IT091 | CAAGCAGAAGACGGCATACGAGAT TGCCAT GTGACTGGAGTTCAGACGTGTGCTCTTC | Fwd primer for 2nd amplification (P7) |

| | | |
|---|---|---|
| oPNL900-Amp2-R-IT055 | AATGATACGGCGACCACCGAGATCTACAC AAGCGA ACACTCTTTCCCTACACGACGCTCTTCCGATCT | Rev primer for 2nd amplification (P5) |
| oPNL901-Amp2-R-IT022 | AATGATACGGCGACCACCGAGATCTACAC CGTACG ACACTCTTTCCCTACACGACGCTCTTCCGATCT | Rev primer for 2nd amplification (P5) |
| oPNL902-Amp2-R-IT084 | AATGATACGGCGACCACCGAGATCTACAC GCACTT ACACTCTTTCCCTACACGACGCTCTTCCGATCT | Rev primer for 2nd amplification (P5) |
| oPNL903-Amp2-R-IT017 | AATGATACGGCGACCACCGAGATCTACAC GTAGAG ACACTCTTTCCCTACACGACGCTCTTCCGATCT | Rev primer for 2nd amplification (P5) |
| oPNL904-Amp2-R-IT095 | AATGATACGGCGACCACCGAGATCTACAC TTCTCC ACACTCTTTCCCTACACGACGCTCTTCCGATCT | Rev primer for 2nd amplification (P5) |
| oPNL926-Amp2-F-IT010 | CAAGCAGAAGACGGCATACGAGAT TAGCTT GTGACTGGAGTTCAGACGTGTGCTCTTC | Fwd primer for 2nd amplification (P7) |
| oPNL927-Amp2-F-IT076 | CAAGCAGAAGACGGCATACGAGAT CGAGAA GTGACTGGAGTTCAGACGTGTGCTCTTC | Fwd primer for 2nd amplification (P7) |
| oPNL928-Amp2-F-IT021 | CAAGCAGAAGACGGCATACGAGAT GTTTCG GTGACTGGAGTTCAGACGTGTGCTCTTC | Fwd primer for 2nd amplification (P7) |
| oPNL929-Amp2-F-IT028 | CAAGCAGAAGACGGCATACGAGAT CAAAAG GTGACTGGAGTTCAGACGTGTGCTCTTC | Fwd primer for 2nd amplification (P7) |
| oPNL930-Amp2-F-IT066 | CAAGCAGAAGACGGCATACGAGAT AGCATC GTGACTGGAGTTCAGACGTGTGCTCTTC | Fwd primer for 2nd amplification (P7) |
| oPNL931-Amp2-R-IT042 | AATGATACGGCGACCACCGAGATCTACAC TAATCG ACACTCTTTCCCTACACGACGCTCTTCCGATCT | Rev primer for 2nd amplification (P5) |
| oPNL932-Amp2-R-IT027 | AATGATACGGCGACCACCGAGATCTACAC ATTCCT ACACTCTTTCCCTACACGACGCTCTTCCGATCT | Rev primer for 2nd amplification (P5) |

| | | |
|---|---|---|
| oPNL933-Amp2-R-IT037 | AATGATACGGCGACCACCGAGATCTACAC CGGAAT ACACTCTTTCCCTACACGACGCTCTTCCGATCT | Rev primer for 2nd amplification (P5) |
| oPNL934-Amp2-R-IT045 | AATGATACGGCGACCACCGAGATCTACAC TCATTC ACACTCTTTCCCTACACGACGCTCTTCCGATCT | Rev primer for 2nd amplification (P5) |
| oPNL935-Amp2-R-IT080 | AATGATACGGCGACCACCGAGATCTACAC GACGGA ACACTCTTTCCCTACACGACGCTCTTCCGATCT | Rev primer for 2nd amplification (P5) |
| *P. fluorescens poly-attB strain construction screening primers* | | |
| oPNL611-3'ampC_fl_F | CTCGGTGAGCAAAACCTTC | flanking primers to screen for insertion of DNA downstream of ampC locus |
| oPNL612-3'ampC_fl_R | TGCTGATGATCGCGATCTA | flanking primers to screen for insertion of DNA downstream of ampC locus |
| oPNL615-3'PFLU5798_fl_F | TGGTTTATGCACTCAACGAG | flanking primers for screen for insertion of DNA downstream of PFLU5798 locus |
| oPNL616-3'PFLU5798_fl_R | ACCCAGACTGACGATGAAG | flanking primers for screen for insertion of DNA downstream of PFLU5798 locus |
| *R. palustris poly-attB strain construction screening primers* | | |
| oPNL413-RPA1300_fl_F | CTTGATGCCCGAAGCCTTCT | primers flanking RPA1300 to screen for its deletion and replacement with poly-attB cassette in Rpalustris CGA009 |
| oPNL414-RPA1300_fl_R | TTCGTCGTCACCTTCGGTTC | primers flanking RPA1300 to screen for its deletion and replacement with poly-attB cassette in Rpalustris CGA009 |
| oPNL415-RPA1300_int_F | CAGTGGGCTGTCGATGTCTT | primers internal to RPA1300 to screen for its deletion in Rpalustris CGA009 |
| oPNL416-RPA1300_int_R | TGAGTTTCGTTTGCACTGCG | primers internal to RPA1300 to screen for its deletion in Rpalustris CGA009 |
| *Primers for screening ts plasmid loss and attP plasmid backbone excision* | | |
| oPNL556 | AGCGAGTCAGTGAGCGA | screening for pGW26/30 plasmid loss |
| oPNL879 | TTCTTAAGATTAACTCACACAGGAGA | screening for pGW26/30 plasmid loss |
| oPNL621 | CATCGGCGTGGTGATATTGG | screening for pGW60 backbone excision by phiC31 integrase |

| | | |
|---|---|---|
| oPNL622 | TGCCGTGGCTGATGATCC | screening for pGW60 backbone excision by phiC31 integrase |
| *Primers for screening attP plasmid insertion in P. fluorescens* | | |
| oPNL622 | TGCCGTGGCTGATGATCC | screening for *attP* plasmid insertion downstream of *ampC* in JE4621 |
| oPNL629 | aaaaccgcccagtctagctatcg | screening of genomic integration of *attP* plasmids |
| *Primers for screening attP plasmid insertion in R. palustris* | | |
| oPNL819 | GTGGCAGAAAGCTTTCACGG | forward primer for screening attR side of pGW60 into ΔRPA_1300::12x polyattB cassette, binds in ColE1/pUC origin |
| oPNL817 | GAGGGGCAGGCAGAACATAG | reverse primer for screening attR side of pGW60 into ΔRPA_1300::10x polyattB cassette, binds in Rpal CGA009 genome |

1 Peabody, G. L., Elmore, J. R., Martinez-Baird, J. & Guss, A. M. Engineered Pseudomonas putida KT2440 co-utilizes galactose and glucose. *Biotechnol Biofuels* **12**, doi:ARTN 295
10.1186/s13068-019-1627-0 (2019).
2 Johnson, C. W. *et al.* Innovative Chemicals and Materials from Bacterial Aromatic Catabolic Pathways. *Joule* **3**, 1523-1537, doi:10.1016/j.joule.2019.05.011 (2019).
3 Quandt, J. & Hynes, M. F. Versatile Suicide Vectors Which Allow Direct Selection for Gene Replacement in Gram-Negative Bacteria. *Gene* **127**, 15-21, doi:Doi 10.1016/0378-1119(93)90611-6 (1993).
4 Yim, S. S. *et al.* Multiplex transcriptional characterizations across diverse bacterial species using cell-free systems. *Mol Syst Biol* **15**, doi:ARTN e8875
DOI 10.15252/msb.20198875 (2019).
5 Bailey, M. J., Lilley, A. K., Thompson, I. P., Rainey, P. B. & Ellis, R. J. Site directed chromosomal marking of a fluorescent pseudomonad isolated from the phytosphere of sugar beet; Stability and potential for marker gene transfer. *Mol Ecol* **4**, 755-763, doi:DOI 10.1111/j.1365-294X.1995.tb00276.x (1995).
6 Larimer, F. W. *et al.* Complete genome sequence of the metabolically versatile photosynthetic bacterium Rhodopseudomonas palustris. *Nat Biotechnol* **22**, 55-61, doi:10.1038/nbt923 (2004).
7 Jayakody, L. N. *et al.* Thermochemical wastewater valorization via enhanced microbial toxicity tolerance. *Energ Environ Sci* **11**, 1625-1638, doi:10.1039/c8ee00460a (2018).
8 Elmore, J. R., Furches, A., Wolff, G. N., Gorday, K. & Guss, A. M. Development of a high efficiency integration system and promoter library for rapid modification of Pseudomonas putida KT2440. *Metab Eng Commun* **5**, 1-8, doi:10.1016/j.meteno.2017.04.001 (2017).
